# Structural basis of biofilm formation mediated by the *Pseudomonas aeruginosa* fibrillar adhesin CdrA

**DOI:** 10.64898/2026.07.13.738186

**Authors:** Olivia E.R. Smith, Camila M. Clemente, Antonina Andreeva, Zhexin Wang, Aaron Weimann, Adam Dinan, Callum Houghton-Flory, Abul K. Tarafder, Jan Böhning, George A. O’Toole, R. Andres Floto, Juan-Carlos Mobarec, Alex Bateman, Tanmay A.M. Bharat

## Abstract

Many bacteria, including the important human pathogen *Pseudomonas aeruginosa*, are naturally found in antibiotic-tolerant, multicellular biofilms. Cell-cell interactions within *P. aeruginosa* biofilms are mediated by a large fibrillar adhesin called CdrA in an extracellular polysaccharide-dependent manner. Here, we report an electron cryomicroscopy structure of the 60 kDa CdrA adhesive N-terminus, which combined with electron cryotomography of focused-ion beam milled specimens, allows us to derive a complete *in situ* model of the native adhesin. Our structure reveals a small adhesive domain (called ADEPT) at the distal tip of CdrA that is nearly perfectly conserved across the *P. aeruginosa* pangenome, with structural similarity to previously reported sugar-binding domains in multiple bacterial species. Inhibitory nanobodies targeting CdrA that reduce biofilm formation bind to epitopes in, or close to, the ADEPT on bacterial cells. Furthermore, structure-guided mutagenesis of residues within the ADEPT abolishes bacterial aggregation, and genomic deletion of the whole ADEPT leads to strong attenuation of biofilm formation. Our data forms a rational basis for future targeted inhibition of pathogenic *P. aeruginosa* biofilms and elucidates the mechanism of biofilm formation mediated by fibrillar adhesins that are widespread in bacteria.

## Introduction

Biofilms are highly organised multicellular communities of bacteria that represent a key mode of prokaryotic life found in nature (O’Toole *et al*., 2000). Within biofilms, sessile bacterial cells are encased in an extracellular matrix (ECM), which is a hallmark of all biofilms, endowing numerous survival advantages to the constituent cells such as tolerance to antibiotic treatment (Flemming *et al*., 2016; Høiby *et al*., 2010). The biofilm ECM is composed of a myriad of polymeric proteins, polysaccharides and extracellular DNA, collectively termed extracellular polymeric substances (Böhning *et al*., 2024; Flemming and Wingender, 2010). In almost all biofilms, bacterial cells express cell-surface adhesins that mediate cell-cell and cell-ECM connections that are crucial to the maintenance of biofilm stability and architecture (Monzon and Bateman, 2022; Smith and Bharat, 2024). Well-characterised examples of such adhesins include RbmA from *Vibrio cholerae* (Giglio *et al*., 2013) and Ag43 from *Escherichia coli* (Heras *et al*., 2014).

*Pseudomonas aeruginosa* is an opportunistic human pathogen that causes chronic infections through the formation of antibiotic-tolerant biofilms (GBD 2021 Antimicrobial Resistance Collaborators, 2024; Thi *et al*., 2020). In biofilms, *P. aeruginosa* expresses the important fibrillar cell-surface adhesin called CdrA, in a cyclic diguanylate (c-di-GMP) dependent manner, to stabilise cell-ECM contacts and mediate biofilm formation (Borlee *et al*., 2010). CdrA is a ∼220 kDa multi-domain protein that shares the overall domain architecture of fibrillar adhesins described previously (Smith and Bharat, 2024; Whelan *et al*., 2021). CdrA has a C-terminal cysteine hook that anchors CdrA to its cognate CdrB pore in the outer membrane, an extension module composed of β-sheet rich repeat domains and an N-terminal adhesion module (Cooley *et al*., 2016; Reichhardt, 2023). Electron cryomicroscopy (cryo-EM) and tomography (cryo-ET) have previously shown that CdrA appears as ∼70 nm matchstick-shaped protrusions extending out from the cell surface (Melia *et al*., 2021). When cytoplasmic c-di-GMP levels are high, CdrA is retained on the bacterial cell surface, promoting cellular tethering and retention within the biofilm through interactions with ECM polysaccharides (Reichhardt *et al*., 2020). At low c-di-GMP levels, the C-terminal retention module of CdrA is proteolytically cleaved by the periplasmic protease LapG, thus shedding CdrA from the cell surface and promoting biofilm dispersion (Cooley *et al*., 2016; Rybtke *et al*., 2015).

CdrA mediates bacterial aggregation by directly binding to the extracellular polysaccharides Psl or Pel through its N-terminal adhesive domain located distal to the cell surface (Melia *et al*., 2021; Reichhardt *et al*., 2020). Psl is composed of repeating units of a neutral pentasaccharide comprising D-glucose, D-mannose and L-rhamnose and is chemically distinct from the cationic N-acetylglucosamine-rich Pel polysaccharide (Byrd *et al*., 2009; Jennings *et al*., 2015). Since Psl and Pel are differentially expressed across *P. aeruginosa* strains, this implies that CdrA has an important role in maintaining biofilm architecture in *P. aeruginosa* (Colvin *et al*., 2012; Weimann *et al*., 2024). In line with this finding, the *cdrA* gene is found in most environmental and clinical isolates characterised (Reichhardt *et al*., 2020), and deletion of CdrA markedly reduces biofilm formation (Borlee *et al*., 2010). Disruption of the CdrA-ECM polysaccharide interaction by addition of monosaccharides such as mannose results in failure of *P. aeruginosa* cells to form stable multicellular aggregates, leading to decreased biofilm biomass (Borlee *et al*., 2010). Furthermore, targeting of CdrA with inhibitory nanobodies renders biofilms more susceptible to antibiotic treatment (Melia *et al*., 2021) and a bispecific monoclonal antibody, MEDI3902, that targets Psl and the type III secretion system has shown promise in animal models and clinical trials for treating *P. aeruginosa* infections (Chastre *et al*., 2022; DiGiandomenico *et al*., 2014; Le *et al*., 2018). Despite the key importance of CdrA, and CdrA-polysaccharide interactions in mediating biofilm architecture, our in-depth understanding of CdrA structure and function remains incomplete. This has limited our ability to exploit CdrA as a potential therapeutic target and has also left gaps in the fundamental knowledge about biofilm formation mediated by fibrillar adhesins.

In this study, we report a cryo-EM structure of the adhesive N-terminal module of CdrA, which showed the presence of a small previously undescribed domain (that we named hereafter ADEPT), which is highly conserved throughout the *P. aeruginosa* pangenome, playing a key role in mediating biofilm formation. We combine our structure with cryo-ET of CdrA on focused ion-beam (FIB)-milled cell surfaces to derive a complete *in situ* model of CdrA, using AlphaFold predictions and molecular dynamics simulations. We further show that inhibitory nanobodies directly bind to CdrA epitopes at or close to the ADEPT and we perform scanning-mutagenesis to map function to our structural data. This study elucidates the mechanism of CdrA-mediated adhesion in biofilms, which will serve as a framework for understanding fibrillar adhesin-mediated biofilm formation in bacteria, assisting in the development of targeted inhibition strategies.

## Results

### Cryo-EM structure of the CdrA adhesive N-terminus

We have previously shown that the tip of CdrA, extending 70 nm away from the *P. aeruginosa* cell surface can be targeted for biofilm inhibition (Melia *et al*., 2021). Since CdrA is anchored to the cell at the C-terminus, we focused our structural studies on the other end of the molecule, at the adhesive N-terminus. For these studies, an N-terminal construct of CdrA was recombinantly expressed and purified (Figs. 1A and S1A-B). The construct consists of the N-terminus of mature CdrA (residues 438-808; thus excluding N-terminal residues 43-437 which are cleaved in mature CdrA (Borlee *et al*., 2010)) and two of the structural repeats comprising the extended stalk (residues 809-989) (Fig. 1A, Methods). The CdrA-438-989 construct was expressed with an N-terminal Maltose Binding Protein (MBP) tag to ensure soluble expression, which was cleaved prior to cryo-EM grid preparation to yield a 60 kDa protein (Fig. 1A). Elongated densities were seen in cryo-EM micrographs, which yielded promising class averages (Figs. 1A, S1C and S1D) showing structural features of the stalk domain, and the matchstick shaped tip, consistent with previous cryo-ET data (Melia *et al*., 2021). We used this dataset to resolve a 3.6 Å-resolution map of the mature CdrA N-terminus (Figs. 1B, S1E-G, S2, Table S1 and Movie S1).

**Fig. 1.**
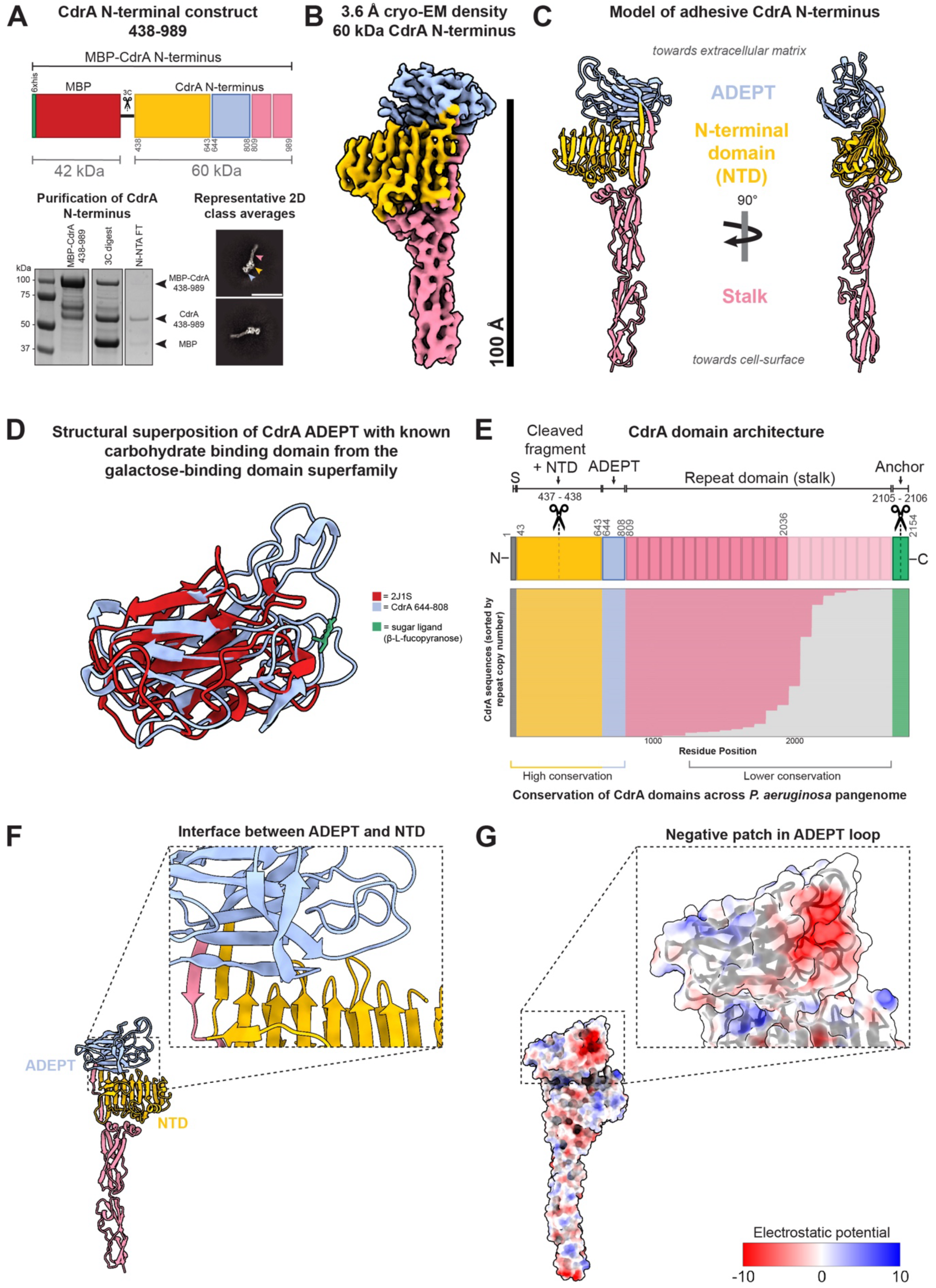
Structure of the adhesive N-terminus of CdrA. **A) Top:** Schematic representation of the CdrA construct consisting of a N-terminal MBP solubility tag, 3C protease site and CdrA residues 438-989. **Bottom:** SDS-PAGE of purification. The MBP-CdrA fusion protein was recombinantly expressed purified by affinity chromatography and gel filtration before MBP cleavage with 3C protease to yield a pure sample of the CdrA-438-989 construct (60 kDa). Ni-NTA = nickel-nitrilotriacetic acid, FT= flow through. Molecular weight standards are indicated on the left. 2D class averages derived from cryo-EM data of the CdrA-438-989 construct. Coloured arrows reference the domains in the schematic representation above. Scale bar represents 150 Å. **B)** A 3.6 Å-resolution map of the 60 kDa CdrA N-terminus. Scale bar indicates 100 Å. **C)** A refined model of the CdrA adhesive N-terminus built into the density showing that the ADEPT (blue) lies distal to the cell surface with the β-strand-rich NTD (yellow) and stalk (pink) below. **D)** Superposition of CdrA ADEPT (blue) with a carbohydrate-binding domain member from the galactose-binding domain superfamily (in red) bound to the sugar β-L-fucopyranose (green) (PDB code: 2J1S). The bound sugar ligand clusters near the loop regions of the CdrA ADEPT. **E) Top:** CdrA domain architecture. Domain boundaries are indicated with numbers. Black = signal sequence (S) (residues 1-42), yellow = N-terminal cleaved fragment + NTD (residues 43-643), blue = ADEPT (residues 644-808), pink = stalk repeats (809-2306), green = C-terminal anchor (2306-2154). The dashed line and scissor symbol between residues 437-438 (separating the N-terminal cleaved fragment + NTD) represents where the N-terminus is cleaved in immature CdrA to release the 43-437 N-terminal fragment (Borlee *et al*., 2010), forming mature CdrA. The scissor symbol between residues 2105-2106 represents where the C-terminal ‘TAAG’ motif can be cleaved by periplasmic protease LapG to release CdrA from the cell-surface (Cooley *et al*., 2016; Rybtke *et al*., 2015). **Bottom:** conservation of CdrA across the *P. aeruginosa* pangenome. Sequences for 998 gapless genomes from RefSeq (Amid *et al*., 2012) where a *bona fide* coding sequence was observed are shown. The adhesive N-terminus (comprising the NTD, ADEPT and first two repeats of the stalk) is near perfectly conserved. Conservation decreases in the stalk, with isolates possessing a different number of stalk repeat regions. CdrA in *P. aeruginosa* strain PAO1 (for which the domain boundaries are shown) has 12 further repeats (opaque pink) whilst other strains can have up to 21 repeats (translucent pink). The C-terminal anchor (green) is well-conserved across the pangenome. **F)** The CdrA N-terminus model shows a tight interface between the ADEPT and NTD. **G)** Electrostatic surface of the CdrA N-terminus, showing a negatively charged patch in the loop region of the ADEPT that is implicated in sugar-binding.

The structure shows clearly resolved secondary structure elements, including β-strands, and side chains (Figs. S1F-G), allowing us to obtain an atomic model of the adhesive tip of CdrA (Fig. 1C). Our model shows that the two stalk domain repeats (pink in Fig. 1C, CdrA residues 809-989) are capped with the N-terminal domain (NTD) of the construct (yellow, CdrA residues 438-643), with a small domain at the distal end (blue, CdrA residues 644-808). Bioinformatic analysis of CdrA revealed that the distal tip domain spanning residues 644-808 belongs to a family of uncharacterised domains that are widely distributed across bacteria and show a propensity to bind to sugars (Figs. 1D and S3A). AlphaFold2 models of members of this family display significant structural similarity to several known carbohydrate binding domains that fold into a galactose-binding, jelly-roll-like β-sandwich fold. Besides adopting this typical fold, these domains are predicted to contain the characteristic structural irregularity in their β-sheet – the so-called β-bulge – that facilitates the binding of sugar molecules and directs its orientation. In CdrA, this domain is a circularly permuted version of the other family members, hence we name it <u>A</u>dhesive <u>D</u>omain <u>E</u>xhibiting <u>P</u>ermuted <u>T</u>opology (ADEPT) (Fig. S3C). Using AlphaFold3 and a peptide sugar mimetic to test *in silico* sugar binding to CdrA and other non-circularly permuted ADEPTs, we found that these domains likely engage sugars through family-conserved residues that define a semi-conserved DSSDY sequence motif (Figs. S3B-D and S7C). Moreover, the predicted sugar binding site is located at the loops, in a mode very similar to that observed in other known, structurally similar sugar binding domains (Figs. S1D, S3A and S3D). Taken together, the cryo-EM structure of CdrA ADEPT and the structural bioinformatics indicate possible modes of CdrA binding to matrix polysaccharides such as Psl.

To further investigate the importance of different parts of CdrA, we analysed CdrA sequences across the *P. aeruginosa* pangenome using 4,289 clinical isolate genomes (Weimann *et al*., 2024) and 1,100 gapless genomes to specifically resolve the repetitive stalk region from RefSeq (Amid *et al*., 2012). Whilst CdrA sequences varied significantly in their stalk domains, particularly in the number of repeats present (3-21 copies), the NTD and the ADEPT show near perfect conservation (Fig. 1E), indicating that these parts of CdrA play a key functional role. Finally, it is noteworthy that the experimental structure of the CdrA adhesive N-terminus (this study) deviates significantly from AlphaFold2 (Jumper *et al*., 2021) predictions (Fig. S4B), which predict an open conformation where the ADEPT is positioned distant from the NTD. Molecular dynamics simulations further indicate that the closed conformation (observed in our cryo-EM structure) is more stable (Figs. S4A-B). Our cryo-EM structure shows that the ADEPT associates tightly with the NTD (Fig. 1F), and exposes a patch of negatively charged residues within a loop region (Fig. 1G).

### Complete architecture of native CdrA using *in situ* cryo-ET

To decipher the organisation of the native, mature CdrA molecule, we utilised a previously reported system of CdrA over-expression, wherein floccules of *P. aeruginosa* could be obtained for cryo-ET analysis (Melia *et al*., 2021). Using an improved workflow for specimen preparation, we applied FIB-milling to obtain lamellae through the CdrA-mediated floccules (Fig. S5A-B). The lamellae were amenable to cryo-ET imaging and allowed us to image native CdrA *in situ*, on the *P. aeruginosa* cell surface (Figs. 2A-C and Movie S2). In reconstructed tomograms, we were able to identify CdrA fibrillar molecules at the cell-surface with a characteristic matchstick-shaped structure, where the N-terminus (NTD + ADEPT) forms the tip (Figs. 2A-C), in line with our previous work. In total, 215 molecules were identified from 23 cells (7 tomograms) at the cell-surface by manual inspection of the tomograms. We measured the length of these molecules manually and found a mean length of 69 nm (SD = 12.7 nm) (Fig. 2D), matching tightly with our previous findings on a smaller dataset. Due to our improved cryo-ET workflows with unfixed cells, improved contrast at the cell surface in tomograms allowed us to observe a diffuse layer of density at the cell surface. We presumed this to be the surface capsule of *P. aeruginosa* cells (Fig. 2C). This putative capsule was found to extend 30 nm from the outer surface of the outer membrane, demonstrating that the adhesive N-terminus of CdrA extends beyond the capsule into the extracellular matrix, where it is accessible for engagement with matrix sugars including Psl or Pel (Figs. 2C-D).

**Fig. 2.**
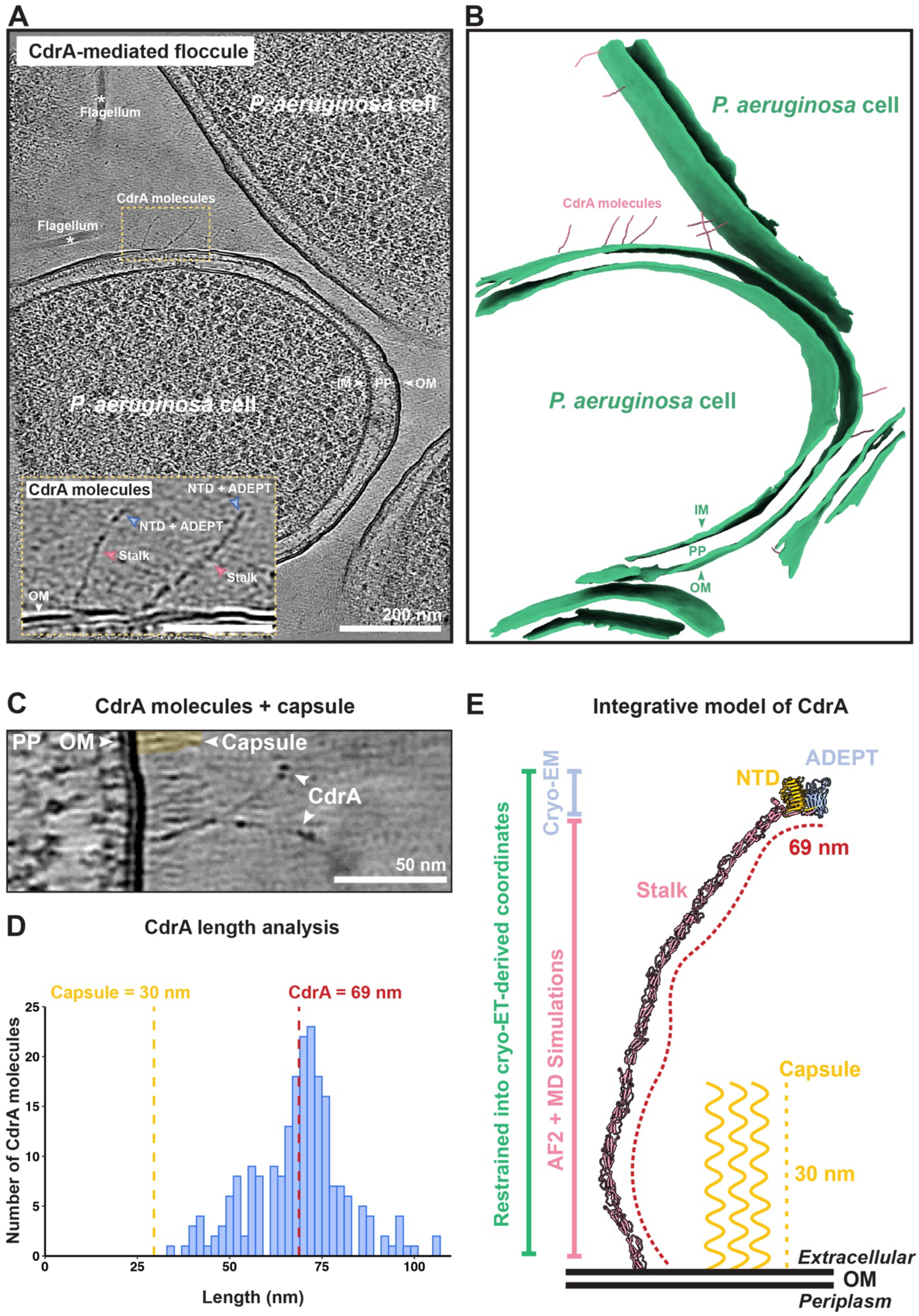
Complete integrative *in situ* model of CdrA at the cell-surface. **A)** Slice through an electron cryotomogram of a FIB-milled CdrA-mediated floccule. Cells and ECM components (e.g. flagella) are visible along with CdrA adhesins (marked by yellow dashed box). IM = inner membrane, PP = periplasm, OM = outer membrane. Scale bar represents 200 nm. The larger yellow dashed box (inset) shows a zoomed subsection of the tomogram (at the area of the smaller yellow dashed box). CdrA molecules are present at the outer membrane, with the NTD and ADEPT distal to the membrane. The stalk of CdrA is also visible. Scale bar of inset image is 70 nm. **B)** Segmentation of the tomogram in panel A shows a number of CdrA molecules (in pink) at the cell surface (in green). Segmentation supports the extended, fibrillar architecture of CdrA. **C)** Zoomed slice through a tomogram showing the cell surface. The capsule (dense layer of cell-surface molecules) is visible, indicated by yellow shading. The capsule extends approximately 30 nm from the cell surface (S.D. = 4.9 nm, n = 232), with CdrA, specifically the NTD and ADEPT extending beyond the capsule. Scale bar represents 50 nm. **D)** Histogram showing the length distribution of manually identified CdrA molecules at the cell-surface. The average length of CdrA was found to be 69 nm (S.D. = 12.7 nm, n = 215) and the capsule thickness 30 nm. **F)** Integrative model of CdrA *in situ* from cryo-ET, single-particle cryo-EM, structure prediction and molecular dynamics (MD) simulations. By combining the cryo-EM structure of the CdrA adhesive N-terminus (Fig. 1) with an AlphaFold2 (AF2) prediction of the remainder of the stalk and restraining this combined full-length CdrA structure into coordinates of CdrA derived from tomograms using molecular dynamics simulations, we were able to create an integrative model of full-length CdrA which is ∼70 nm in length. The capsule is shown in yellow with respect to CdrA. OM = outer membrane.

Using the *in situ* cryo-ET data, we next sought to obtain a full-length structural model of CdrA, by combining our experimental data with computational approaches. To this end, we connected our experimental cryo-EM structure of the CdrA N-terminus with an AlphaFold2 (Jumper *et al*., 2021) prediction of the remaining stalk domain sequence from our *P. aeruginosa* PAO1 strain. This remaining stalk domain sequence was modelled in overlapping constructs using AlphaFold2 and assembled along the cryo-ET-derived molecular coordinates (Figs. S5C-D) using Modeller (Webb and Sali, 2016). This initial integrated model was subsequently refined by molecular dynamics (MD) simulations performed in explicit solvent, yielding a full-length structural model of CdrA that was consistent with the *in situ* cryo-ET data (Fig. 2E).

### Mapping nanobody binding sites to functional inhibition of CdrA

Having produced a full-length model of CdrA, we next investigated why different CdrA-binding nanobodies from our original panel (Melia *et al*., 2021) had a range of inhibitory effects and whether these activities could be explained by nanobody binding to different epitopes within CdrA. We selected three nanobodies which disrupted CdrA-mediated floccules to different levels, including two which also had differing inhibitory activity against *P. aeruginosa* biofilms (Melia *et al*., 2021). Our previously reported Nb_CdrA_ (henceforth referred to as Nb6) was found to be the most disruptive against bacterial flocculation, while Nb2 was the least disruptive and Nb7 showed an intermediate level of disruption (Fig. 3A).

**Fig. 3.**
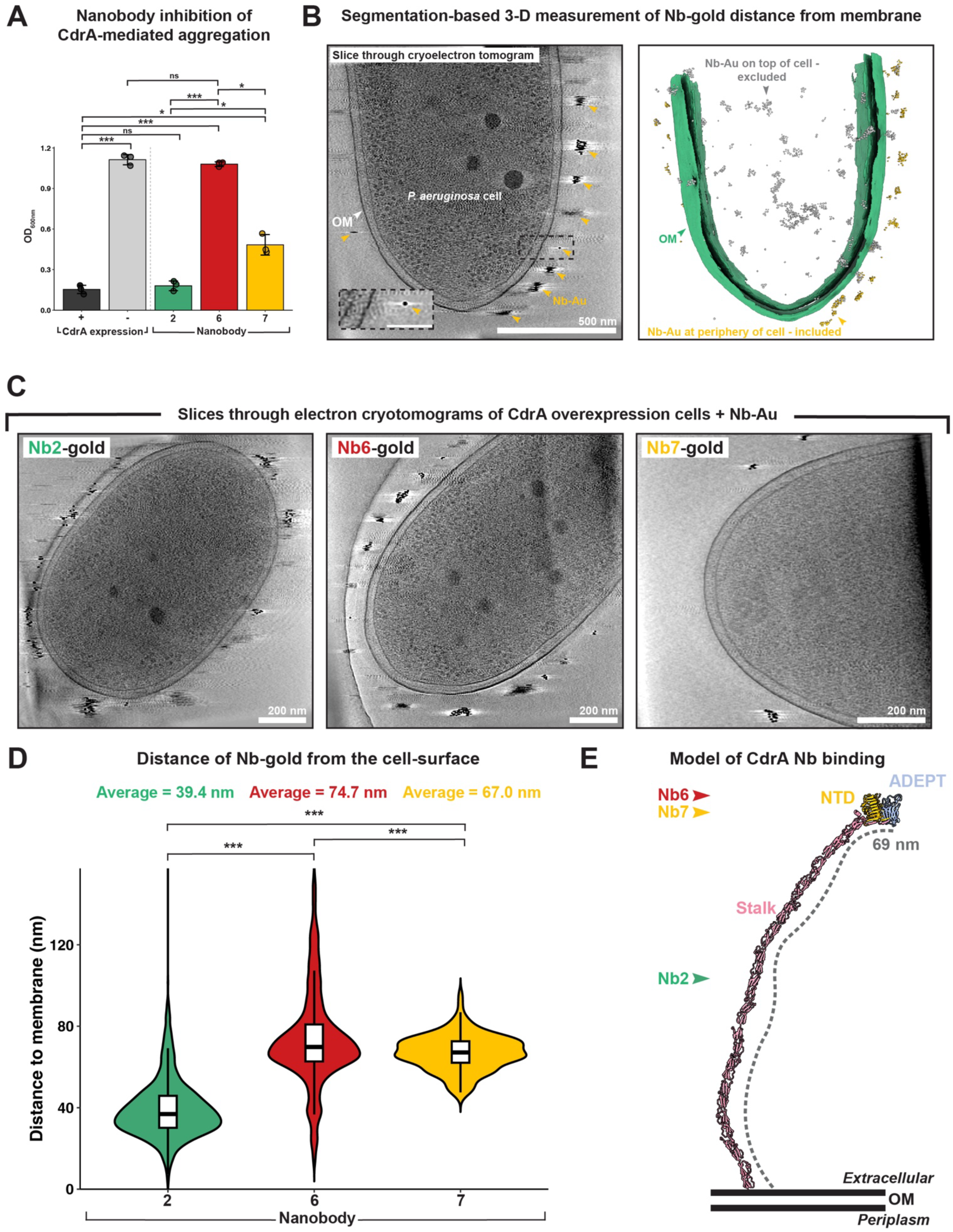
Mapping nanobody binding sites to functional inhibition of CdrA. **A)** Bar chart displaying the inhibitory activity of three anti-CdrA nanobodies, assayed using the CdrA-overexpression flocculation system. CdrA-mediated floccules (biofilm-like aggregates in solution) have a low optical density at 600 nm (OD_600_, dark grey bar). Conversely, without CdrA, bacteria cannot flocculate and form a standard turbid bacterial solution with a high OD_600_ (light grey bar). The same assay showed that Nb2 did not inhibit CdrA-mediated floccules (low OD_600_ values) (green bar), Nb6 was a potent inhibitor of CdrA-mediated flocculation (high OD_600_ values) (red bar) and Nb7 showed an intermediate level of inhibition (yellow bar). The assay was performed in triplicate with error bars indicating standard deviation. Significance symbols represent Holm-adjusted p-values for Welch’s t test between the groups specified by horizontal bars, where: n.s. = p > 0.05, * = p ≤ 0.05, ** = p ≤ 0.01, *** = p ≤ 0.001. **B)** Slice through electron cryotomogram (left hand image) showing gold-labelled nanobodies at the cell-surface (indicated by gold arrow heads). Zoomed inset shows an individual Nb-gold at the cell surface. Segmentation of Nb-gold labelled cell (right hand image) shows membrane (green) and Nb-gold. For quantification of Nb-gold distance, Nb-gold on top of the cell (in grey) was excluded from measurements due to inaccuracy of measuring distance to the membrane. Nb-gold at the periphery of cells (in yellow) was retained for measurements. **C)** Slices through electron cryotomograms for each of the Nb-gold conjugates (Nb-Au) show high levels of surface decoration for each Nb-Au. Scale bars are 200 nm. **D)** Violin plot displaying the distribution of 3D distances of each of the Nb-gold from the cell surface. Measurements of Nb-gold were conducted on three high-quality tomograms of cells for each Nb. For the boxplot, the horizontal central line represents the median, the box represents the interquartile range and the whiskers represent observations within 1.5x the interquartile range. Non-inhibitory Nb2-Au was located closest to the membrane, with an average distance of ∼39 nm. Inhibitory Nb6-Au was located the farthest from the cell-surface, on average ∼75 nm from the cell surface. Nb7-Au was found on average 67 nm from the cell-surface. Significance symbols represent Holm-adjusted p-values for Welch’s t test between the groups specified by horizontal bars, where: *** = p ≤ 0.001. **E)** Plotting of Nb-gold distance from the membrane onto integrative model of CdrA. When plotted on to our integrative model of CdrA (see Fig. 1), we observe that the most inhibitory Nb6 targets the adhesive N-terminus of CdrA, most likely the ADEPT. Nb2 targets the stalk and Nb7 most likely targets the NTD. Binding locations of each nanobody are represented by arrowheads with labels.

To relate these inhibitory activities to nanobody binding sites on CdrA, we conducted epitope mapping using cryo-ET. All three nanobodies were first conjugated with gold nanoparticles and were then incubated with cells over-expressing CdrA, and control cells with no CdrA expression (Fig. S6). After washing to remove unbound nanobody, cells expressing CdrA were found to be highly decorated with gold nanoparticles (Fig. 3B-C). In contrast, the cell surface of CdrA deletion controls lacked this surface decoration (Fig. S6). We measured the distance of gold particles from the *P. aeruginosa* outer membrane in three dimensions (3D), through segmentation of gold and the outer membrane, showing the average location of each nanobody along the CdrA molecule (Fig. 3B). The least disruptive nanobody, Nb2, was bound ∼39 nm away from the cell surface, corresponding to a location on the CdrA stalk domains (Fig. 3C-E), illustrating that binding to the stalk had no effect on the ability of CdrA to mediate biofilm formation. Nb7 was bound ∼67 nm from the cell surface, placing it near the NTD and ADEPT, while gold nanoparticles conjugated with Nb6 were positioned ∼75 nm away from the cell surface, close to the tip of CdrA (Fig. 3C-E). This nanobody mediated epitope mapping indicates that the most disruptive nanobody is likely directly engaging with the ADEPT at the tip of CdrA molecules on the native *P. aeruginosa* cell surface. This probably disrupts the ability of CdrA to bind to Psl and other matrix polysaccharides, further highlighting the importance of the ADEPT in biofilm formation.

### Deletion of the ADEPT and ADEPT:NTD interface residues diminishes cell-cell interactions

Since inhibitory nanobodies all bound close to the ADEPT, we next wanted to probe the importance of the ADEPT in biofilm formation. First, we analysed publicly available transcriptomic datasets of planktonic *P. aeruginosa* compared with bacteria from biofilms (Thöming *et al*., 2020), which confirmed that both *cdrA* and *cdrB* genes are upregulated in biofilms (Fig. 4A). Next, we tested wild-type bacteria (WT), and isogenic genomically mutated strains with cdrA deleted completely (Δ*cdrA*) or with solely the ADEPT deleted (Δ*ADEPT*) for their ability to form biofilms. In a static biofilm assay (O’Toole, 2011), the Δ*cdrA* strain showed strongly attenuated biofilm formation compared to the WT as determined by crystal violet staining of bacterial biomass, as expected from previous reports (Borlee *et al*., 2010) (Fig. 4B). In line with our data above, the Δ*ADEPT* also showed biofilm attenuation, nearly to the level of the Δ*cdrA* strain (no significant difference with the Δ*cdrA* strain), confirming the importance of this domain in biofilm formation (Fig. 4B).

**Fig. 4.**
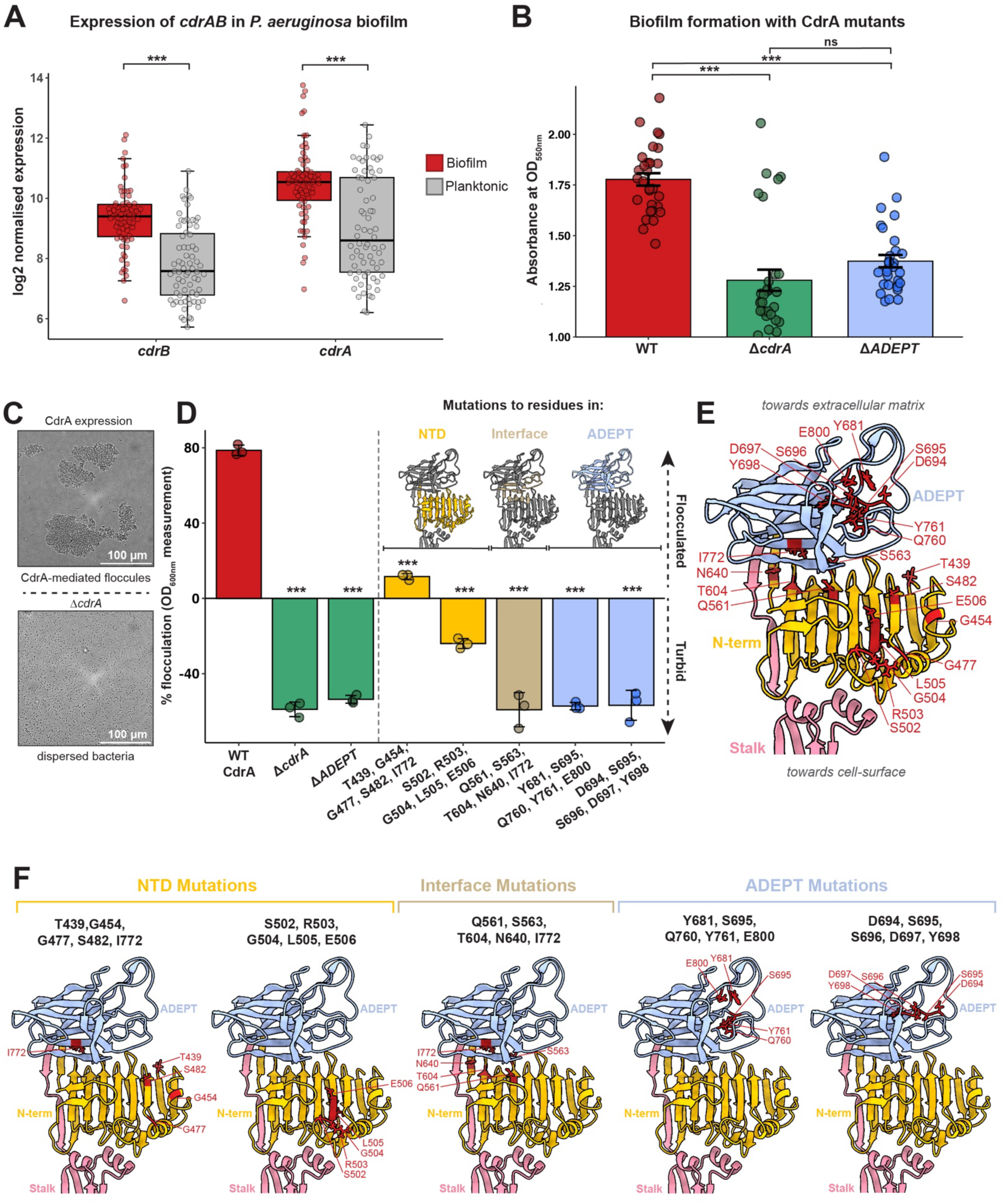
Disruption of CdrA:sugar binding. **A**) Transcriptomic analysis shows that both *cdrA* and *cdrB* (cognate pore which holds CdrA in the outer membrane) are transcriptionally upregulated in biofilms (red bar) compared to planktonic cells (grey bar). Boxplots show the median value (central line), the interquartile range (box) and the whiskers extend to values within 1.5x the interquartile range. Benjamini-Hochberg adjusted *p*-values (for comparisons between biofilm and planktonic) are: *cdrA* – 5.24 x 10^-6^, *cdrB* – 1.08 x 10^-11^. On the plot, p-values are represented by: *** = p ≤ 0.001. **B)** Quantification of biofilm biomass from genomic mutants of *P. aeruginosa* in a static biofilm assay. Deletion of genomic *cdrA* (green bar) or solely the *ADEPT* (blue bar) attenuates biofilm formation to similar levels, as quantified by crystal violet staining. WT biofilm biomass is indicated by a red bar. Black overlaid error bars show standard error of the mean. The assay included 10 technical replicates in triplicate and significance symbols represent Holm-adjusted p-values for Welch’s t test between the groups specified by horizontal bars, where: ns = p > 0.05, and *** = p ≤ 0.001 **C)** Brightfield images of CdrA-mediated floccules (top image) and disrupted CdrA-mediated floccules (bottom image). Scale bars represent 100 µm. **D)** Quantification of the ability of the bacteria to form floccules after mutation of the putative sugar-binding residues. Flocculation can be quantified using OD_600_ measurements, with floccules having a low OD_600_ and dispersed bacteria having a high OD_600_. Deletion of the entire ADEPT decreased flocculation to a similar level observed for deletion of CdrA (green bars). Mutations to the ADEPT (blue bars) and ADEPT:NTD interface (beige bar) were found to reduce flocculation to a level similar to deletion of the entire ADEPT (green bar). NTD mutations (yellow bars) were found to reduce flocculation but not to the same extent as ADEPT or ADEPT:NTD interface mutations. The flocculation assay was performed in triplicate and error bars represent standard deviation. Significance symbols represent Holm-adjusted p-values for Welch’s t-test between each WT CdrA and each mutant group, where: *** = p ≤ 0.001. See Table S2 for sources of each mutation set. **E)** Mutated residues mapped on the cryo-EM structure of the adhesive N-terminus (CdrA-438-989). Mutated residues are shown in red with their side chains atoms displayed and labelled. **F)** Mutations are also shown mapped individually to the cryo-EM structure of the adhesive N-terminus based on the mutation set from which they came.

Next, to interrogate the role of putative sugar-binding residues within the ADEPT and NTD, we conducted *in silico* alanine scanning mutagenesis based on our structure. We then used free energy perturbations in MD simulations to observe how each alanine mutation altered free energy changes involved with Psl binding (compared with the respective non-mutated residue) (Fig. S7A). These simulations highlighted several residues in the ADEPT, the NTD and in the interface of ADEPT:NTD that could potentially reduce the ability of CdrA to bind to polysaccharides (Table S2). We also selected several putative sugar-binding residues based on conservation seen in the ADEPT family from AF3 sugar-binding predictions (Fig. S3D, Table S2) and the motif conservation (Fig. S7C, Table S2). Furthermore, we also identified putative NTD sugar-binding residues through structural alignment of a right-handed β-helix domain, member of the same pectate lyase superfamily as the CdrA NTD (Fig. S7B). To test the effect of mutating these residues to alanine experimentally, we again turned to our CdrA over-expression system, where mutant proteins could be conveniently expressed at the *P. aeruginosa* cell surface, and the effect of the mutation followed by testing the ability of *P. aeruginosa* expressing mutant CdrA to mediate flocculation (Fig. 4C).

For each mutant, five residues were mutated to alanine and expression of the penta-alanine mutant CdrA at the cell surface confirmed by shearing cell surface molecules and assaying them on a protein gel. All mutants expressed a CdrA molecule of the expected molecular weight at the cell surface (Fig. S7E). Next, we tested the ability of each mutant to mediate flocculation in our *in vitro* flocculation assay. As controls, we found that compared to WT (genomic deletion of *cdrA* complemented with *cdrA* on an inducible plasmid), Δ*cdrA* (genomic deletion of *cdrA* with no *cdrA* on a plasmid) showed an extremely attenuated ability to flocculate (Fig. 4C-D). In line with our biofilm crystal violet assay, Δ*ADEPT* showed attenuated ability to flocculate, nearly as weak as the Δ*cdrA* strain. Next, mutations in the NTD also showed attenuated cell-cell interactions (Figs. 4C-D and S7), but the effect was not as strong as Δ*ADEPT*. Finally, mutations in either the ADEPT:NTD interface, or of the putative sugar binding residues in ADEPT showed strong attenuation of flocculation, nearly to the same level as Δ*ADEPT or* Δ*cdrA*. These results together indicate that residues in the loop region of ADEPT (Fig. 1, 4E-F and S7), and those in the ADEPT:NTD interface are key to the ability of CdrA to aggregate, presumably through sugar (Psl) binding (Fig. S7D). Our results establish the CdrA-ADEPT as a key mediator of cellular aggregation and biofilm formation in *P. aeruginosa*, showing how a small protein domain in a large fibrillar adhesin can mediate multicellular community formation.

## Discussion

Adhesion is a critical process in biofilm formation and the maintenance of biofilm architecture (Böhning *et al*., 2023; Schluter *et al*., 2015) and fibrillar adhesins play a key role in this process (Monzon and Bateman, 2022; Smith and Bharat, 2024). In this study, we present a complete *in situ* model of the critical fibrillar adhesin, CdrA, which mediates cell-ECM contacts in *P. aeruginosa* biofilms by binding extracellular polysaccharides. We combined bioinformatic analyses, cryo-EM structures, *in situ* cryo-ET, with molecular dynamics simulations and mutagenesis experiments to delineate the importance of CdrA, and putative sugar binding residues of CdrA, in biofilm formation.

Our structural and bioinformatic analysis of CdrA identified a previously uncharacterised domain within its N-terminus – the ADEPT. The ADEPT forms a β-sandwich fold with similarity to the galactose-binding domain superfamily and is widely distributed in many compositionally diverse proteins involved in adhesion across bacterial species (Fig. S3). In CdrA, the ADEPT exhibits circular permutation of the domain topology compared to other members of the family, highlighting the structural plasticity and adaptability of this fold. Proteins in this family show high levels of sequence diversity, which may underpin substrate specificity and avidity. Furthermore, relative orientations and conformational dynamics of this domain family are poorly predicted, emphasising the need for experimental structure determination. In stark contrast to the sequence diversity within the family, the ADEPT is almost perfectly conserved across the *P. aeruginosa* pangenome, highlighting its functional importance in this pathogen.

Interestingly, our experimentally determined structure of the CdrA N-terminus shows the ADEPT tightly interacting with the NTD in a ‘closed’ confirmation (Fig. 1), whereas AlphaFold3 modelling predicts a more ‘open’ conformation (Fig. S4B). This mismatch could be suggestive of a clamping mechanism of polysaccharide binding between these two domains. Indeed, structure-based mutagenesis of residues in both the ADEPT and NTD disrupted floccule formation in our *in vitro* assays (Fig. 4). The involvement of two domains in polysaccharide binding may increase the avidity of the interaction or broaden the substrate specificity. Some ADEPT-like-domain-containing proteins have either tandem copies of the ADEPT-like domain or additional adhesive domains, suggesting that this may be a common theme amongst polysaccharide binding proteins. Nevertheless, mutagenesis of the ADEPT had the greatest effect on flocculation and a CdrA Δ*ADEPT* mutant showed diminished biofilm formation, suggesting that the ADEPT is the key mediator of cell-ECM interactions (Fig. 5).

**Fig. 5.**
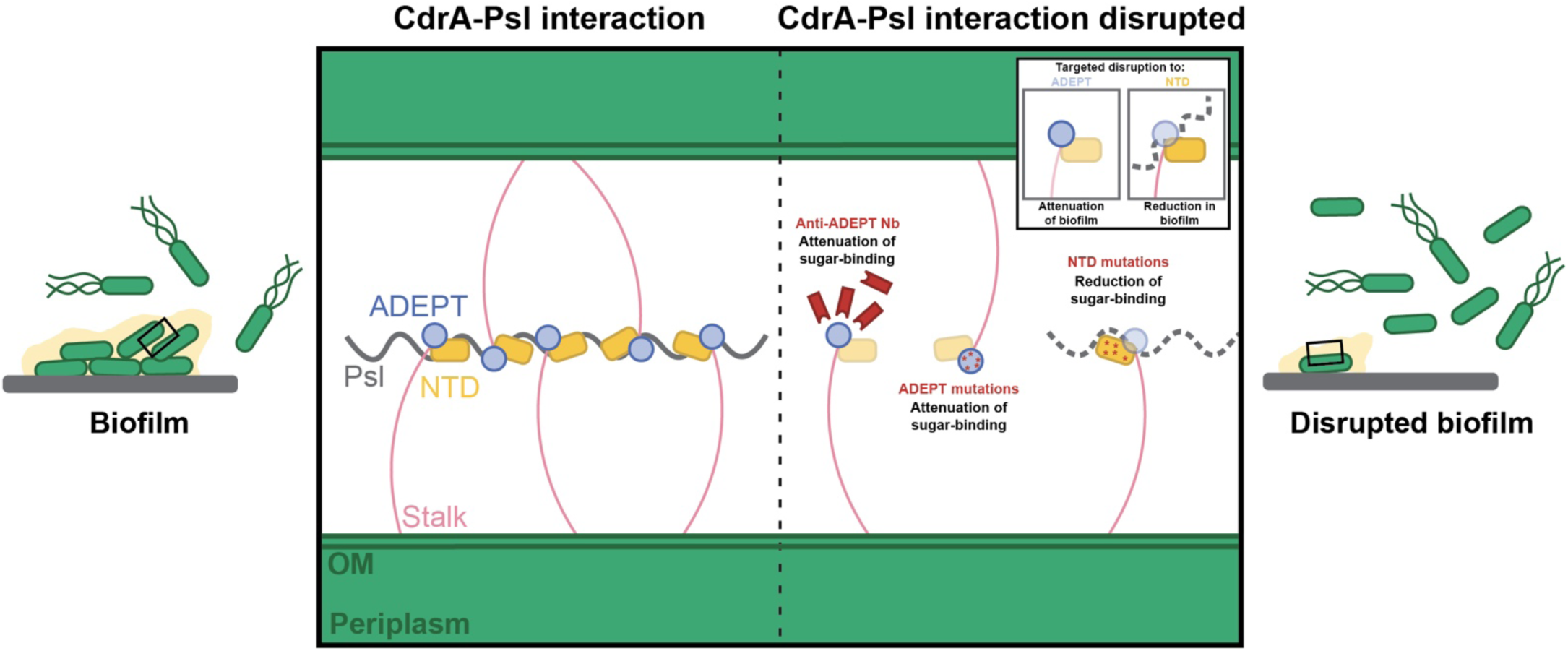
The ADEPT is crucial to CdrA function in *P. aeruginosa* biofilm formation. Schematic showing the importance of the ADEPT and NTD of CdrA in *P. aeruginosa* biofilm formation. In the biofilm state, CdrA adheres to the polysaccharide Psl predominantly through interactions with the ADEPT and the ADEPT:NTD interface. Disruption of the ADEPT-Psl or ADEPT:NTD-Psl interaction through mutation or inhibitory nanobodies results in biofilm attenuation. Disruption to the NTD results in biofilm reduction but not to the same extent as ADEPT disruption.

Our cryo-EM, *in situ* cryo-ET and MD simulation-derived, complete model of CdrA shows the position of the ADEPT relative to the cell (Fig. 2). This is a powerful demonstration of how cryo-ET can contribute to the understanding of pathogenic molecules in their native state (Clemente *et al*., 2026). Our model shows an extended conformation, where the stalk allows the adhesive tip to clear the bacterial capsule and reach extracellular polysaccharide within the biofilm matrix. The repeat regions themselves show the greatest variation in sequence and copy number in the pangenome (Fig. 1), consistent with the idea that CdrA functions as a periscope protein (Whelan *et al*., 2021), where modulation of repeat number possibly tunes stalk length to accommodate different capsule thicknesses or distant ECM sugar components.

Targeting CdrA-mediated adhesion is a promising strategy for treating biofilm infections, and we previously showed that nanobodies targeting CdrA can disrupt *P. aeruginosa* biofilms (Melia *et al*., 2021). By combining our full-length model of CdrA with cryo-ET of gold labelled nanobodies, we could map nanobody epitopes and correlate binding site to functional inhibition, revealing that nanobodies binding to the CdrA N-terminus, close to the ADEPT, were most effective at disrupting CdrA function and impairing biofilm formation. This finding could form the basis for the development of more targeted therapeutic strategies against *P. aeruginosa* biofilms.

In summary, we have provided a comprehensive *in situ* structural model of the *P. aeruginosa* adhesin CdrA, filling a critical gap in our mechanistic and structural understanding of biofilm formation mediated by this molecule. We identified and structurally characterised the ADEPT and showed it is a key determinant in binding extracellular polysaccharides to maintain biofilm architecture. Our results are not only of interest from a biomedical perspective but also illuminate how fibrillar adhesins could mediate biofilm formation in other environmental and pathogenic bacteria.

## Materials and Methods

### Expression and purification of the CdrA N-terminus (CdrA-438-989)

A synthetic GeneBlock (Thermo Fisher Scientific) containing the *E. coli* codon optimised DNA sequence of CdrA from residues 438-989 was used, which was flanked with restriction sites for NcoI and XhoI. Both the pETM44 plasmid (Addgene) and the synthetic GeneBlock were digested with NcoI and XhoI for 1 hour (New England Biolabs). Each respective digested fragment was isolated using gel extraction (Qiagen) and were ligated to produce pETM44-CdrA-438-989. pETM44-CdrA-438-989 was transformed into chemically competent BL21 (DE3) cells by heat shock. Starter cultures of the cells were prepared and used to inoculate 24 l of Luria broth (LB) at a dilution of 1:40. Cultures were grown at 37 °C with agitation at 180 revolutions per minute (rpm) until reaching an OD_600_ (optical density at 600 nm wavelength of light) of 0.5. Protein expression was induced with the addition of 1 mM isopropyl β-D-thiogalactoside (IPTG), with cultures being incubated overnight at 18 °C with shaking at 180 rpm. Cultures were centrifuged at 4000x*g* for 20 minutes and cell pellets stored at –80 °C. Pellets were resuspended in lysis buffer (50 mM Tris-HCl pH 7.5, 500 mM NaCl, 0.5 mM TCEP, 5 mM MgCl_2_, DNAse I, cOmplete EDTA-free protease inhibitor (Roche)) and were disrupted using an initial sonication step followed by disruption at 30 kpsi using a cell disruptor (Constant Systems). The MBP-CdrA fusion protein was applied to a nickel-nitrilotriacetic acid (Ni-NTA) column equilibrated in Ni-NTA buffer (50 mM Tris-HCl pH 7.5, 500 mM NaCl, 0.5 mM TCEP, 20 mM imidazole) and was eluted with Ni-NTA buffer with 500 mM imidazole. Fractions containing MBP-CdrA fusion protein were combined and applied to a HiLoad 16/600 Superdex 200 pg column pre-equilibrated in buffer (50 mM Tris-HCl pH 7.5, 500 mM NaCl, 0.5 mM TCEP). After gel filtration, MBP-CdrA-438-989 was incubated with 3C protease overnight at 4 °C to facilitate MBP cleavage. The cleaved MBP tag and hexahistidine-tagged 3C protease were removed using Ni-NTA resin to yield a pure sample of CdrA-438-989 (60 kDa).

### Sample preparation for cryo-EM

For single particle cryo-EM, purified MBP-CdrA-438-989 or CdrA-438-989 (MBP tag cleaved) was diluted with PBS to a resultant concentration of 3 µM. Next, 2.5 µl of the diluted sample was applied to freshly glow-discharged Quanitfoil R2/2 Cu/Rh 200 mesh grids. Grids were plunge-frozen using a Vitrobot mark IV (Thermo Fisher Scientific) following a 60 second wait time with a blot time of 2.5 seconds.

### Cryo-EM data collection

For the three MBP-CdrA-438-989 datasets and one of the CdrA-438-989 (MBP cleaved) datasets, single particle cryo-EM data was collected using a Titan Krios G3 microscope (Thermo Fisher Scientific) operating at 300 kV equipped with a Selectris energy filter (slit width 20 eV) and a Falcon 4i direct electron detector (Thermo Fisher Scientific). Data was collected at a nominal magnification of 130,000 x (with a resultant pixel size of 0.955 Å/pixel) with a defocus range of – 1.25 µm to –2.5 µm running in counting super-resolution mode. Across the three datasets, a total of 11,655 movies were collected without a stage tilt and 3,512 movies were collected with a 30° stage tilt using EPU (Thermo Fisher Scientific), each with a total dose of 40 e^-^/Å^2^, with each movie consisting of 40 frames. For the remaining CdrA-438-989 (MBP cleaved) dataset, data was collected using a Titan Krios G3 microscope (Thermo Fisher Scientific) operating at 300 kV equipped with a Quantum energy filter (slit width 20 eV) and a K3 direct electron detector (Gatan). Data was collected at a nominal magnification of 105,000 x (with a resultant pixel size of 0.825 Å/pixel) with a defocus range of –1.25 µm to –2.5 µm in counting super-resolution mode. A total of 29,439 movies were collected using EPU (Thermo Fisher Scientific), with a total dose of 40 e^-^/Å^2^, respectively, with each movie consisting of 40 frames.

### Cryo-EM data processing

Data processing was performed in CryoSPARC v4.5.1 (Punjani *et al*., 2017). *Preprocessing*: For each of the five datasets, movies were first patch-based motion corrected and subjected to patch-based CTF estimation. Micrographs were filtered and excluded based on ice quality and CTF-fit resolution. Micrographs were then denoised using the cryoSPARC micrograph denoiser tool to aid in particle picking. *Exploratory data processing and generation of a model for particle picking*: To generate a template for picking datasets, the cryoSPARC blob picker (with a target particle size of 150 to 250 Å) was used to pick particles on a subset of the data (0.955 Å/pixel dataset). A total of 1,938,538 particles were picked. The picked particles were extracted with a 320 pixels^2^ box size and Fourier-cropped to 120 pixels^2^, and subjected to several 2D classification rounds, with particles belonging to well-aligned 2D class averages of CdrA-438-989, being selected. After 2D classification, 127,048 particles remained. These particles were re-extracted in a 320 pixels^2^ box size (with no Fourier crop to the box size). Particles were then subjected to *ab initio* reconstruction and rounds of heterogeneous refinement to further clean the particle stack. Non-uniform refinement (Punjani *et al*., 2020) of the resulting 119,351 particles yielded a volume with features consistent for those expected with the filamentous architecture of CdrA. *Building stacks of good particles for merging datasets*: For each of the datasets, a low-pass-filtered version of this volume was used to generate templates which were then low-pass filtered to 20 Å to pick particles. After template picking, particles were extracted in a box size of 424 pixels^2^ (corresponding to ∼405 Å for the 0.955 Å/pix datasets) and were downsampled by a factor of 4 and subjected to a round of 2D classification. The particle set was gently cleaned, removing obvious junk only (e.g. edges), and then subjected to heterogeneous refinement against 5 decoy classes and the low-pass filtered CdrA-438-989 class. This classification in 3D was repeated several times. For the large dataset of CdrA-438-989 (29,439 movies), top views and side views were classified separately to prevent loss of the rarer top views of CdrA-438-989. *Combining datasets*: To merge datasets of different pixel sizes, the boxscaler.py script (https://github.com/maxewilkinson/CryoEM-scripts) was used to determine a box size (with the constraint of requiring a box size of approximately 300 Å to accommodate double the longest axis of CdrA-438-989) and Fourier crop for the 0.85 Å/pix dataset which resulted in a pixel size of 0.95 Å/pix. To this end, particles belonging to the 0.85 Å/pix dataset were re-extracted in a box of 382 pixels^2^ with a Fourier crop of 330 pixels^2^ whilst particles belong to the 4 datasets collected at a pixel size of 0.95 Å were re-extracted in a 330 pixels^2^ box (without a Fourier crop). After merging, the combined 565,313 particles were subjected to heterogeneous refinement against 5 decoy classes and a low-pass-filtered volume of CdrA-438-989 (see steps above). Non-uniform refinement was performed on particles in the good class (which was always the most populous class) to monitor how particle cleaning affected resolution. The new volume from non-uniform refinement was used as the low-pass filtered good class for subsequent heterogeneous refinement. These iterations were repeated until there were very few particles being classified into the junk classes (achieved after approximately 10 rounds of this classification). After this particle cleaning, 104,619 particles remained. These particles were subjected to local CTF refinement and then to a consensus refinement with the cryoSPARC non-uniform refinement which yielded a 3.44 Å resolution reconstruction using the gold-standard 0.143 FSC criterion. Local refinement with a mask which excluded the most C-terminal repeat (which looked flexible and less well-resolved in previous refinements), resulted in a map with a slightly worse estimate of 3.6 Å resolution. However, a manual inspection of the map from the local refinement showed better resolved side chain densities and separation of the β-strands in the NTD and ADEPT.

### Data visualisation, analysis and model building

For model building, the AlphaFold3 prediction of the CdrA 438-989 adhesive N-terminus was used (Abramson *et al*., 2024). This included an additional 4 N-terminal residues (sequence: GPMG) which were present after 3C cleavage of the N-terminal MBP tag. The AlphaFold3 prediction was split into 3 domains: NTD, ADEPT and stalk were separately rigid body-fitted into the density in UCSF ChimeraX (Meng *et al*., 2023). ISOLDE (Croll, 2018) was used to improve the fit of each domain into the density before inspection in Coot (Emsley *et al*., 2010). In Coot, the fit was inspected and side chains positions were modified where there was visible side chain density. The model was subjected to several rounds of refinement using PHENIX (Liebschner *et al*., 2019), ISOLDE (Croll, 2018) and manual correction in Coot (Emsley *et al*., 2010). Model validation was performed in PHENIX (Liebschner *et al*., 2019) and data visualisation was performed in UCSF ChimeraX (Meng *et al*., 2023). After building of the model into the density, for display of the model in figures, the scar residues from the 3C cleavage (sequence: GPMG) were removed so that the N-terminus of the model starts with CdrA residue 438. Furthermore, for clarity, all residue numbers provided in figures are in reference to full-length CdrA residues as opposed to residue numbers in the CdrA-438-989 construct. Cryo-EM images or cryo-ET slices were prepared using IMOD (Kremer *et al*., 1996) and Fiji (Schindelin *et al*., 2012).

### Bioinformatic identification of CdrA ADEPT

The ADEPT was initially identified as a putative adhesive domain from the AlphaFold model of the *Pseudomonas aeruginosa* UniProtKB:Q9HVG6 protein between residues 644 to 808. Using the HMMER package, homologues were collected by iteratively searching the UniProt Reference Proteome sequences above a threshold of 27 bits. Similarity was found to UniProtKB:A0A1U7I4B5 and UniProtKB:A0A2G4F0Q5 strongly suggestive of homology, which was supported by the conservation of the most conserved motifs. The position of these motifs, however, was different suggesting a circular permutation. Sugar binding prediction was performed using AlphaFold server (Abramson *et al*., 2024) using a tetra-threonine peptide glycosylated with mannose and glucose in first, second and fourth position and producing models with ipTM: 0.8 and pTM: 0.79.

### Pangenome analysis of CdrA

Clinical isolate genome assemblies (Weimann *et al*., 2024) were searched for the CdrA ADEPT and NTD using tblastn (Altschul *et al*., 1997) using the *P. aeruginosa PAO1* sequence. As next-generation sequenced-based assemblies are known to fail to accurately resolve repeats (Treangen and Salzberg, 2011), gapless genome assemblies were downloaded from RefSeq (Amid *et al*., 2012) to characterise the repeat stalk region of CdrA. The repeat region was anchored by searching each gapless genome for the CdrA C and N-terminal regions using tblastn (Altschul *et al*., 1997). Only complete protein sequences, i.e. those without stop codons or truncations, were retained for visualisation.

### Molecular dynamics simulations of CdrA

MD simulations of CdrA were performed to assess the conformational stability of the AlphaFold2 (AF2) predicted model. The AF2 model (generated using AlphaFold2 (Jumper *et al*., 2021), top-ranked by pLDDT, mean pLDDT = 89.04) corresponding to residues (438-908) was used as the starting structure for MD simulations. The model was prepared using the Protein Preparation Wizard, and the prepared structure was then used as input for the Desmond System Builder. Systems were built by solvating the protein in an orthorhombic SPC water box with a 30 Å buffer in each dimension and neutralizing with Na^+^ ions; the OPLS4 force field was applied throughout. Production MD was run in Desmond (NPT ensemble, 300.0 K, 1.01325 bar) following the default relaxation protocol, with trajectories recorded every 500 ps and energies every 1.2 ps. Three independent 500 ns replicas were run. Trajectory analysis was carried out using the Desmond Trajectory Analysis tools. The whole process was performed using the Schrödinger Suite (release 2025-3).

### Cryo FIB milling and tomography of CdrA floccules

An overnight culture of *P. aeruginosa* PAO1 Δ*cdrA* pMQ72-*cdrAB* was subcultured in LB media and grown to OD_600_ of 0.9 and induced with 2% arabinose for 1 hour until floccules could be directly observed. 2.5 µl of the floccule suspension was applied to gold grids with R1/4 holey SiO_2_ film (Quantifoil). Excess liquid was blotted from the backside of the grid for 15 s before plunge freezing using a Vitrobot (Thermo Fisher Scientific). Frozen grids were clipped in FIB-specific AutoGrids (Thermo Fisher Scientific) before transfer into a Crossbeam FIB/SEM dual-beam microscope (Zeiss) equipped with a cryo stage (Quorum). The grids were first sputter-coated with platinum followed by another coating of metalloorganic platinum through a gas-injection-system. The floccules were milled into lamellae of ∼150 nm following a four-step milling strategy with decreasing FIB currents: 700 pA, 300 pA, 100 pA and 50 pA. The milled lamellae were transferred into a Titan Krios (Thermo Fisher Scientific) equipped with a K3 BioQuantum detector with an energy filter (Gatan). Tomographic datasets were acquired using SerialEM (Mastronarde, 2003) at 42,000x nominal magnification (pixel size 2.13 Å) tilting from –54° to +54° with 3° increment. Tomograms were reconstructed using AreTomo (Zheng *et al*., 2022) after being pre-processed in Warp (Tegunov and Cramer, 2019). For visualisation, tomograms were denoised using CryoCARE (Buchholz *et al*., 2019) and low-pass filtered in EMAN2 (Tang *et al*., 2007).

### Measurement and visualisation of cell-surface CdrA molecules

CdrA molecules were manually annotated in IMOD (Kremer *et al*., 1996). CdrA molecules were picked in the plane which best showed their density and were identified according to the criteria that they extended from the membrane, possessed a length >40 nm and a consistent density across the length. Interpolation of picked points was done using the AddModPts function from IMOD, with the interpoint distance being 4.26 nm. Radius of curvature was also calculated for the points. For visualisation, cell membrane was segmented using Membrain (Lamm *et al*., 2022) and annotated CdrA molecules were converted to cylindrical volumes. Images were rendered in ChimeraX (Meng *et al*., 2023). For the estimation of capsule size, up to five capsule measurements per z-plane were manually recorded based on the observed extent of increased contrast variation around the membrane for each cell compared to the bulk intercellular space, with measurements repeated every 20-30 Z-planes (17-26 nm). It was noted that this layer was not visualised around all cells in the dataset and was dependent on tomogram quality.

### Construction of the integrated model of CdrA

From the measurement of cryo-ET CdrA length data, three-dimensional coordinates of the CdrA fibrillar adhesin were extracted from cryoelectron tomograms. For the adhesive N-terminus (comprising the NTD, ADEPT and two stalk repeats), the cryo-EM structure was used directly. The remaining sequence of the protein was divided into overlapping constructs spanning the remainder of the stalk domain, each predicted independently using AlphaFold2 (Jumper *et al*., 2021); all predictions yielded a global pLDDT score above 90. Both the cryo-EM structure and the AlphaFold2 stalk models were assembled into a full-length model using Modeller (Webb and Sali, 2016), with the cryo-ET-derived coordinates serving as a spatial template to constrain the overall conformation. The assembled model was subjected to molecular dynamics simulations using the Desmond module (Schrödinger Release 2025-3) with the OPLS4 force field in an orthorhombic box with explicit SPC water. An initial equilibration stage was performed with the protein fully restrained to allow solvent relaxation. Production simulations of 50 ns were then carried out at 300 K and 1.0325 bar with harmonic positional restraints applied to the backbone, allowing local structural relaxation while preserving the global conformation consistent with the cryo-ET-derived coordinates.

### Nanobody expression and purification

Nanobodies against CdrA were raised in a previous study (Melia *et al*., 2021). To express nanobodies, nanobody phagemids were first transformed by heat shock into chemically competent WK6 *E. coli* cells. Overnight cultures of WK6 cells containing nanobody phagemids were subcultured into 12 l of Terrific Broth (TB). Cultures were grown at 37°C until mid-log phase (OD_600_ was used to assess growth) and then induced with IPTG and incubated overnight with shaking at 180 rpm at 20 °C. The following morning, cells were pelleted by centrifugation at 4,000 x *g* for 20 minutes and frozen. Frozen cell pellets from the 12 l of culture were resuspended in 180 ml TES buffer (200 mM Tris-HCl pH 8.0, 0.5 mM EDTA, 500 mM sucrose) and subjected to osmotic shock (by addition of 360 ml of a 1 in 4 dilution of the TES buffer) to lyse the periplasm. Hexahistidine-tagged nanobodies were applied to a Ni-NTA column pre-equilibrated in Ni-NTA buffer (50 mM Tris-HCl pH 7.4, 500 mM NaCl, 0.5 mM TCEP, 20 mM imidazole) before being eluted with Ni-NTA buffer supplemented with 500 mM imidazole. Further purification was achieved by application of the nanobody-containing fractions to a Superdex 75 Increase 10/300 GL column pre-equilbrated in gel filtration buffer (50 mM Tris pH 7.4, 150 mM NaCl, 0.5 mM TCEP).

### Nanobody-mediated disruption assays

A culture of *P. aeruginosa* PAO1 Δ*cdrA* pMQ72-CdrAB (arabinose-inducible, CdrA overexpression strain) (Melia *et al*., 2021) was prepared in LB supplemented with 30 µg/ml gentamicin and grown overnight at 37 °C with shaking at 180 rpm. The following morning, the overnight cultures were normalised to an OD_600_ of 1. Next, 100 ml of LB (supplemented with 30 µg/ml gentamicin) containing 2.5% of the normalised overnight culture and 2% arabinose were incubated at 37 °C with shaking at 180 rpm for 3.5 hours. 3 ml of culture was aliquoted and incubated with 60 µl of 250 µM nanobody (resulting in a final nanobody concentration of 5 µM). After an incubation period of 2 hours, 1 ml of culture was added to cuvettes and the OD_600_ was recorded.

### Nanobody-gold conjugation and labelling

Nanobody fractions were diluted to 10 µM and subsequently dialysed overnight into 20 mM Tris-HCl pH 7.5, 150 mM NaCl in preparation for gold conjugation. Dialysed nanobody was incubated with a 20-molar excess of Ni-NTA-5 nm gold (Nanoprobes) for 30 minutes at room temperature. Overnight cultures of *P. aeruginosa* PAO1 Δ*cdrA* pMQ72-*cdrAB* (arabinose-inducible, CdrA overexpression strain) and *P. aeruginosa* PAO1 Δ*cdrA* pMQ72 (control) in LB supplemented with 30 µg/ml gentamicin were produced. Overnight cultures were sub-cultured 1:30 in LB (supplemented with 30 µg/ml gentamicin) and grown for a total of 6 hours at 37 °C with shaking at 180 rpm, with 2% arabinose added to induce CdrA expression after 2 hours. After 6 hours, 0.5% mannose was added to the cultures to disrupt floccules with gentle agitation and cultures were normalised to an OD_600_ of 0.4. 100 µl of cell suspension and 100 µl of nanobody solution were incubated together (producing a 0.1 µM resultant concentration of nanobody-gold) for 30 minutes at room temperature. Cells were pelleted by centrifugation at 4000x*g* for 10 minutes and the pellet was resuspended in 200 µl of PBS. This wash step was repeated a further time remove any unbound nanobody-gold or unbound gold before a final centrifugation step. The final pellet was resuspended in 25 µl of PBS ahead of cryo-EM grid preparation.

### Cryo-ET of nanobody-labelled cells

For all the nanobody-labelled datasets, 2.5 µl of the washed cells were applied to freshly glow-discharged Quantifoil R2/2 Cu/Rh 200 mesh grids. Grids were plunge frozen using a Vitrobot Mark IV (Thermo Fisher Scientific) following a wait time of 5 s with a blot force of –10 and a blot time of 2 s. For Nb6-gold and Nb2-gold datasets, tilt series were acquired on a Titan Krios microscope operating at 300 kV with a Selectris energy filter (slit width 20 eV) and Falcon 4i direct electron detector (Thermo Fisher Scientific). Tilt series were acquired with SerialEM (Mastronarde, 2003) at a nominal magnification of 42,000 x (with a calibrated pixel size of 3.01 Å/pix) with a total dose of 120 e/Å^2^ over the entire tilt series. The tilt series were collected between –60° and +60° with a 3° tilt increment. Three high quality tilt series were collected for each Nb-gold with a defocus range between –4 to –6 µm. For the Nb7-gold dataset, tilt series were acquired on a Titan Krios microscope operating at 300 kV with a Quantum energy filter (slit width 20 eV) and K3 direct electron detector (Gatan). Tilt series were acquired with SerialEM (Mastronarde, 2003) at a nominal magnification of 42,000 x (with a calibrated pixel size of 2.13 Å/pix) with a total dose of 121 e/Å^2^ across the entire tilt series. The tilt series were collected between –60° and +60° with a 3° tilt increment. 3 tilt series were collected with a defocus range between –4 to –6 µm. All tomograms were reconstructed using the Relion-5 tomography pipeline (Burt *et al*., 2024), with manual correction of the gold fiducial model performed in IMOD (Kremer *et al*., 1996). Tomograms were visualised in IMOD and Fiji (Kremer *et al*., 1996; Schindelin *et al*., 2012).

### Nanobody-gold distance measurement

Cell membranes were segmented using a self-trained 2.5D U-Net implemented in Dragonfly (Comet Technologies). Gold particles were segmented using thresholding and separated into instances by connection. The centre of mass of each instance was considered as the position of each gold particle. The gold particles that are on the top or the bottom of cells were not considered for measurement as the membrane segmentation in those regions was not available due to the missing wedge artefact. Distance between gold particles and cell membrane was calculated using a customised script, which created a 3D gold distance map for every tomogram based on the membrane segmentation.

### Transcriptomic analysis of CdrA

RNA-Seq data from Thöming *et al*. (Thöming *et al*., 2020) were quantified by pseudo-alignment to strain-specific gene sets using Kallisto (Bray *et al*., 2016). Strain-specific gene sequences were derived from the pangenome analysis performed in Weimann *et al*. (Weimann *et al*., 2024). Differential expression analysis was performed using DESeq2 (Love *et al*., 2014). For visualization of gene expression, raw read counts were scaled using library-specific scaling factors from DESeq2 and log2-transformed.

### Construction of genomic Δ*ADEPT*

The genomic Δ*ADEPT* strain was constructed by allelic exchange following previously reported methods (Hmelo *et al*., 2015). Briefly, a GeneBlock (Thermo Fisher Scientific) of the DNA sequence 400 bp upstream of the ADEPT, the DNA sequence of a 5x GS linker (to replace the ADEPT and flanking β-strands between CdrA residues 641 and 811) and the 400 bp region downstream of the ADEPT was constructed. The GeneBlock was PCR amplified and flanked with a BamHI and HindIII sites which facilitated ligation into a BamHI and HindIII digested pEXG2 suicide vector. pEXG2-delADEPT was transformed into *E. coli* Jke201 (donor strain) by heatshock and positive clones were confirmed by PCR. 500 µl of an overnight culture of *E. coli* Jke201 pEXG2-delADEPT grown in LB supplemented with 100 µM diaminopimelic acid (DAP) was mixed with 500 µl of an overnight culture of *P. aeruginosa* PAO1. Cells were pelleted at 9000x*g* and the pellet was resuspended in 1 ml of fresh LB. The centrifugation step was repeated once more and the pellet resuspended in 100 µl of fresh LB. To facilitate conjugation, the resuspended pellet was spotted on a 25 mm filter paper disc (with 0.45 µm pore size) on a LB agar plate supplemented with 100 µM DAP. The plate was incubated for 6 hours at 37 °C after which, the filter paper was removed and was vortexed in 1 ml of LB to resuspend cells. The resuspended cells were plated on a LB agar plate containing 30 µg/ml gentamicin and incubated at 37 °C for 24 hours. Colonies were picked from the plate and grown in aliquots of 2 ml of LB supplemented with 30 µg/ml gentamicin for 4 hours. Counter-selection was achieved by streaking each culture on a no salt LB agar plate containing 10% sucrose. Colonies were screened for successful introduction of the Δ*ADEPT* mutation using PCR and subsequently confirmed using whole genome sequencing.

### Static biofilm assay

Crystal violet quantification of static biofilm mass was performed as described previously (O’Toole, 2011). Briefly, overnight cultures of PAO1 WT, PAO1 Δ*cdrA* and PAO1 Δ*ADEPT* were sub-cultured to a resultant dilution of 1 in 20 in M9 media. Cultures were grown for 3 hours at 37°C with shaking at 180 rpm. All cultures were normalised to an OD_600_ of 0.3 before 200 µl of each culture was pipetted into each well of a flat-bottomed 96-well plate (Corning). The plate was grown statically at 37°C for 24 hours to cultivate biofilm. After 24 hours, the cultures were tipped out of wells and the plate was washed in distilled water, excess water was removed by tapping the inverted plate on paper towels. A total of 3 washes were performed before 200 µl of 0.1% crystal violet solution was added to each well. After a 15-minute incubation with crystal violet, the wells were subjected to 3 washes (with the same method as above). Plates were left inverted overnight to dry. 200 µl of 30% acetic acid was added to wells and incubated for 15 minutes. The contents of each of the wells was then transferred to a new 96-well plate and crystal violet staining was quantified by reading the plate at OD_550_.

### Selection of putative sugar-binding residues for mutation

*Free energy perturbation calculation:* To identify residues important for Psl recognition, we performed free energy perturbation (FEP) residue scanning across three putative binding sites on CdrA (ADEPT, ADEPT:NTD interface, and NTD). Psl was prepared using LigPrep, including conformational sampling, and CdrA was prepared using the Protein Preparation Wizard. Psl was then docked onto each of the three sites using Glide, with the docking grid centred on each site and sized according to the dimensions of Psl. The top-ranked pose by GlideScore at each site was carried forward for FEP residue scanning. For each site, every residue within 4 Å of the bound ligand was individually mutated to alanine using the Residue Scanning module of FEP+, yielding ΔΔG values that delineate the relative energetic contribution of each residue to Psl recognition. FEP+ calculations were run using the OPLS4 force field in the µVT ensemble, with a 0.02 ns equilibration step followed by 10 ns of production simulation for each stage of the thermodynamic cycle, using the default sampling scheme (12 λ-windows). Systems were solvated with the SPC water model. All calculations were performed using Schrödinger Release 2025-3 (Schrödinger, LLC). *Structural alignment with PDB:2PYH:* Putative NTD sugar-binding residues were identified by superimposing the NTD domain with the protein PDB: 2PYH, a member of the same superfamily (IPR006633), which mapped the known 2PYH sugar-binding site onto the CdrA NTD and defined the docking grid used for subsequent Psl docking (Glide, Schrödinger Release 2025-3, Schrödinger, LLC). Residues contacting the docked Psl and based on their location within a region of notable sequence conservation in the CdrA NTD-2PYH alignment were selected for mutagenesis.

### Expression of mutant CdrA molecules in a CdrA-overexpression system

A synthetic GeneBlock (Thermo Fisher Scientific) was designed to contain the *cdrAB* operon with internal, silent restriction sites for MauBI and BsiWI located before the NTD and after the ADEPT respectively (between residues 396 and 868). The GeneBlock was flanked with EcoRI and HindIII restriction sites to facilitate ligation into pMQ72 (CdrAB expression plasmid). The GeneBlock and pMQ72 were digested with EcoRI and HindIII (New England Biolabs) for 1 hour and were run on a 1% agarose gel to separate digested fragments. Fragments of interest were gel excised and purified (Qiagen) and were ligated to make pMQ72-CdrAB-derivative. Subsequently, smaller GeneBlocks corresponding to the residues between MauBI and BsiWI (including the NTD and ADEPT) were designed for each of the CdrA mutants. For all mutants, sets of 5 residues identified either from FEP experiments (see above) or from bioinformatic comparisons were mutated to alanine residues. A total of 3 FEP mutant GeneBlocks and 2 bioinformatic-based mutants were constructed. All mutant GeneBlocks and the recipient pMQ72-CdrAB-derivative were digested with MauBi and BsiWI (Thermo Fisher Scientific), gel excised and fragments of interest were ligated, producing pMQ72-CdrAB-mutant. Correct insertions were confirmed by plasmid sequencing.

### Quantification of ability of mutant CdrA to flocculate

Overnight cultures were prepared for each of the *P. aeruginosa* PAO1 pMQ72-CdrAB-mutant strains in LB supplemented with 30 µg/ml gentamicin and were grown at 37 °C with shaking at 180 rpm. The following morning, overnight cultures were normalised to an OD_600nm_ of 2. Cultures of 100 ml LB (supplemented with 30 µg/ml gentamicin) with 2.5% normalised overnight culture were grown for 2.5 hours at 37 °C with shaking at 180 rpm. The OD_600_ was recorded, then 2% arabinose was added to induce mutant CdrA expression and cultures were allowed to grow for a further 3.5 hours. The OD_600_ was finally recorded as a measurement of flocculation. To ensure that all mutant CdrA constructs were still expressed at the cell surface, 50 ml aliquots of each of the cultures were centrifuged at 4,000x*g* for 20 minutes to pellet cells. The pellets were resuspended in 250 µl of phosphate-buffered saline (PBS) and were vortexed for 90 seconds to shear mutant CdrA from the cell surface. One ml of the sheared material was centrifuged at 21,000x*g* for 40 minutes to pellet cell debris. An aliquot of the supernatant was concentrated in a 0.5 ml 10 kDa molecular weight cut-off Amicon spin concentrator before both the supernatant (unconcentrated) and the concentrated supernatant were applied to an SDS-PAGE gel to check for the presence of mutant CdrA.

## Acknowledgements

This work was supported by the Medical Research Council, as part of United Kingdom Research and Innovation (also known as UK Research and Innovation) [Programme MC_UP_1201/31 to T.A.M.B]. This work was supported through a research collaboration between AstraZeneca UK Limited and the UK Medical Research Council (Blue Sky Programme – Project BSF2-11). For the purpose of open access, the MRC Laboratory of Molecular Biology has applied a CC BY public copyright license to any Author Accepted Manuscript version arising. T.A.M.B. would like to thank the Wellcome Trust (grant 225317/Z/22/Z) and the Lister Institute for Preventative Medicine for support. R.A.F., A.W. and A.D. were supported by a Wellcome Discovery award (226602/Z/22/Z) and a grant from the LifeArc/Cystic Fibrosis Trust Innovation Hub (THUB01) for CF Lung Health and Infection. G.A.O was supported by funding from National Institutes of Health (NIH/R37-AI83256). We thank Ray Owens and Jiandong Huo for raising some of the nanobodies from the panel reported previously and providing plasmids encoding nanobodies. We would also like to thank the MRC LMB electron microscopy, biophysics and scientific computing facilities. We also thank Diamond for access and support of the cryo-EM facilities at the UK national electron Bio-Imaging Centre (eBIC), proposal BI37660.

## Data and Materials Availability

Further information and requests for resources and reagents should be directed to and will be fulfilled by Tanmay Bharat. The nanobodies described in this paper, and where applicable their sequences, may be made available by the MRC Laboratory of Molecular Biology upon request under the terms of an MTA with restrictions on commercial use, confidentiality and publication.

## Data and code availability

All relevant data is presented in the manuscript, and there is no specific novel code associated with this study.

## Author contributions

OERS, CMC, AA, ZW, AW, AD, CHF, AKT, JB, JCM, AB, TAMB were responsible for resource provision, investigation and data validation. OERS, CMC, AA, ZW, AW, AD, CHF, JB, JCM, AB, TAMB were responsible for data visualisation. GAOT, RAF, JCM, AB and TAMB co-ordinated the project administration, supervision and the acquisition of funding. OERS, AKT and TAMB were responsible for the conceptualisation and preparation of the original draft. All co-authors reviewed and edited the manuscript.

## Supplementary figures

**Fig. S1.**
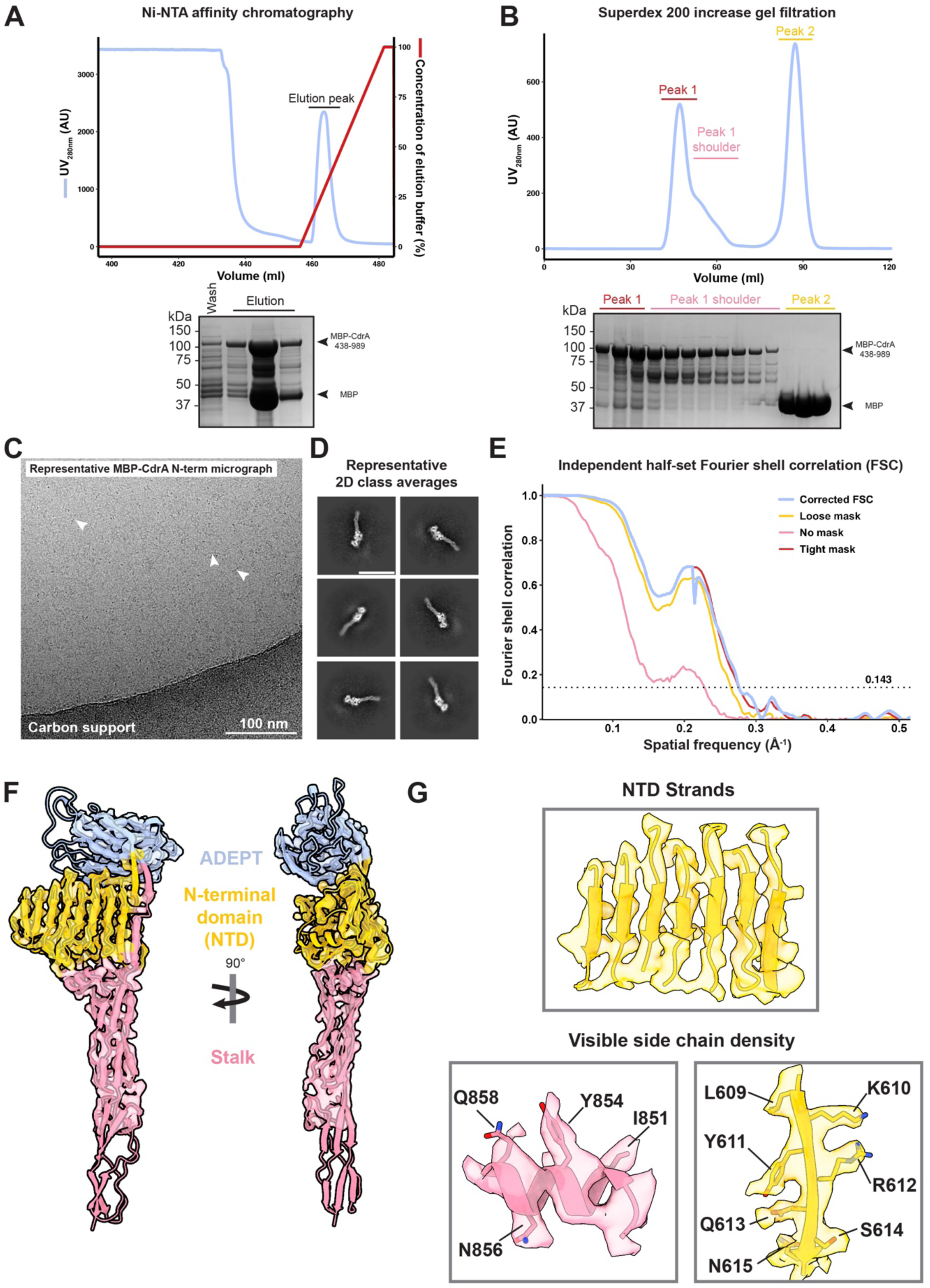
Single particle cryo-EM structure of the CdrA adhesive N-terminus. **A)** Nickel immobilised metal ion affinity chromatography of MBP-CdrA-438-989 construct. Chromatogram (top) displays a large UV peak in the elution which corresponded to the MBP-CdrA-438-989 construct when analysed by SDS-PAGE (bottom). Molecular weight standards are indicated on the left. The construct was further verified by peptide fingerprinting mass spectrometry. **B)** Gel filtration of the affinity purified MBP-CdrA-438-989 yielded a series of peaks in the UV trace (top). SDS-PAGE analysis (bottom) shows that peak 1 shoulder corresponds to MBP-CdrA-438-989 and was used for downstream work. Peak1 corresponds to soluble aggregates due to their proximity to the void volume. Peak 2 corresponds to free MBP. **C)** Representative micrograph of grids prepared with purified MBP-CdrA-438-989. White arrow heads point to representative particles. **D)** Representative 2D class averages obtained from averaging the merged single particle cryo-EM datasets of MBP-CdrA-438-989 and CdrA-438-989 (MBP-cleaved version). Secondary structure features are apparent in all parts of the protein. Scale bar = 150 Å. **E)** Fourier shell correlation (FSC – blue line) as calculated by cryoSPARC for the local refinement job which was used to produce the density for model building. The 0.143 criterion used for resolution estimation is shown as a dotted line. Masked curves are shown in different colours. Resolution of map = 3.62 Å. **F)** Fit of CdrA adhesive N-terminus model into the cryo-EM map. The mask used in local refinement excluded the terminal repeat hence the absence of strong density for this region. Domains are coloured, with ADEPT = blue, NTD = yellow, stalk = pink. **G)** Zoom in of different parts of the model into map fit. Top: fit of the model in the sheet region in the NTD. Residues shown in cartoon schematic (from left to right β-strands: 439-445, 484-496, 512-519, 534-541, 565-572, 586-594, 606-614). Bottom left: side chain density for residues in the stalk helix: I851, Y854, N856 and Q858. Bottom right: side chain density for residues in one of the NTD strands: L609, K610, Y611, R612, Q613, S614, N615. Residue numbers refer to residues in full-length CdrA. Side chains are labelled by heteroatom (with nitrogen atoms represented in blue and oxygen atoms in red) and hydrogen atoms have been hidden.

**Fig. S2.**
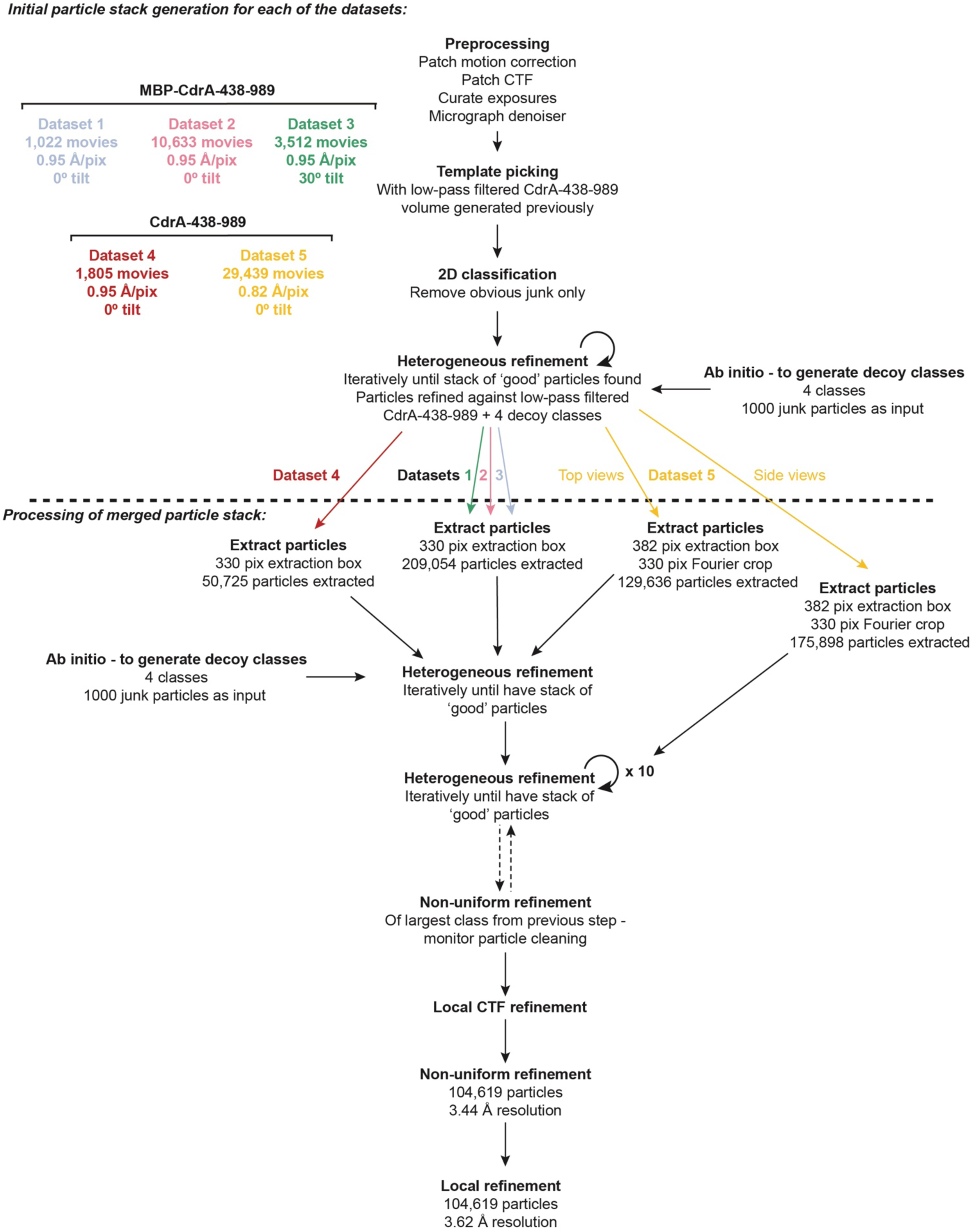
Simplified single particle cryo-EM workflow for CdrA adhesive N-terminus structure determination. Data processing workflow of individual datasets (top) to produce a stack of particles which was merged (bottom) and particles processed together further to yield a volume at 3.62 Å resolution (as calculated by cryoSPARC using the 0.143 GS-FSC criterion). All processing steps were carried out using cryoSPARC v4.5.1.

**Fig. S3.**
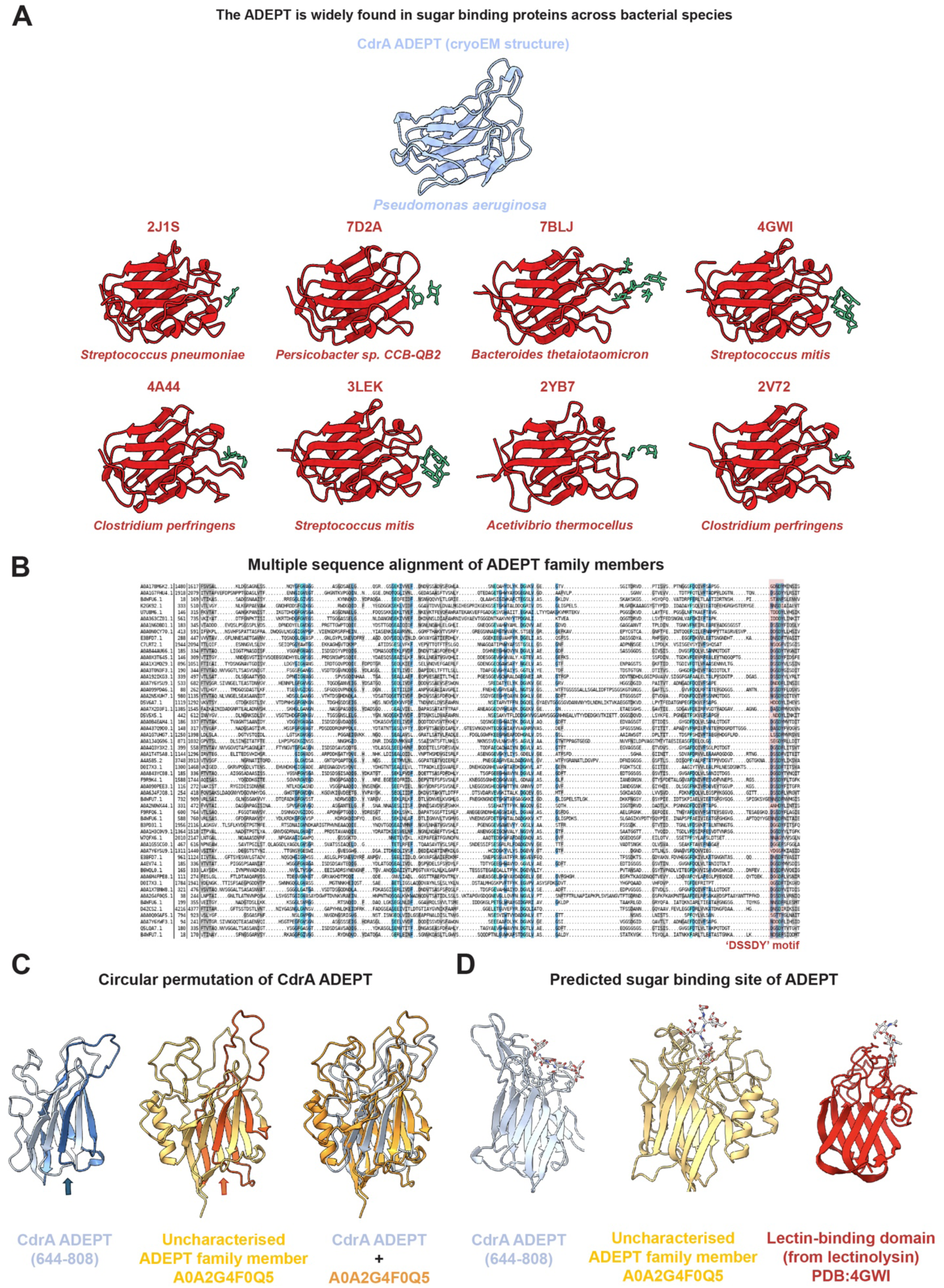
Circular permutation in CdrA ADEPT and similarity of its predicted sugar binding site to known adhesive domains. **A)** Cryo-EM structure of the CdrA ADEPT (residues 644 – 808) with the experimental structures of other structurally-similar domains, members of the galactose-binding domain superfamily (coloured red; Protein Data Bank (PDB) codes provided) with their co-crystallised sugar ligands (green). **B)** Multiple sequence alignment of ADEPT family members. Colours in MSA represent conservation of residues and proteins are listed with their Uniprot codes. The location of the conserved ‘DSSDY’ motif is highlighted in red. **C)** Comparison of the AlphaFold2 (AF2) models of the CdrA ADEPT (residues 644 – 808, shown in blue) and an uncharacterised member of ADEPT family (UniprotKB:A0A2G4F0Q5, residues 1 – 250, shown in orange). The circularly permuted region, which is located at the N-terminus of CdrA ADEPT and at the C-terminus of A0A2G4F0Q5 ADEPT, is shown in dark blue and red-orange, respectively (with coloured arrows to indicate this). **D)** Side-by-side comparison of the AlphaFold3 (AF3) predicted models of CdrA (shown in blue) and A0A2G4F0Q5 (shown in orange) bound to peptide sugar mimetic; and the crystal structure of the lectin binding domain of lectinolysin (shown in red) in complex with Lewis y (PDB:4GWI).

**Fig. S4.**
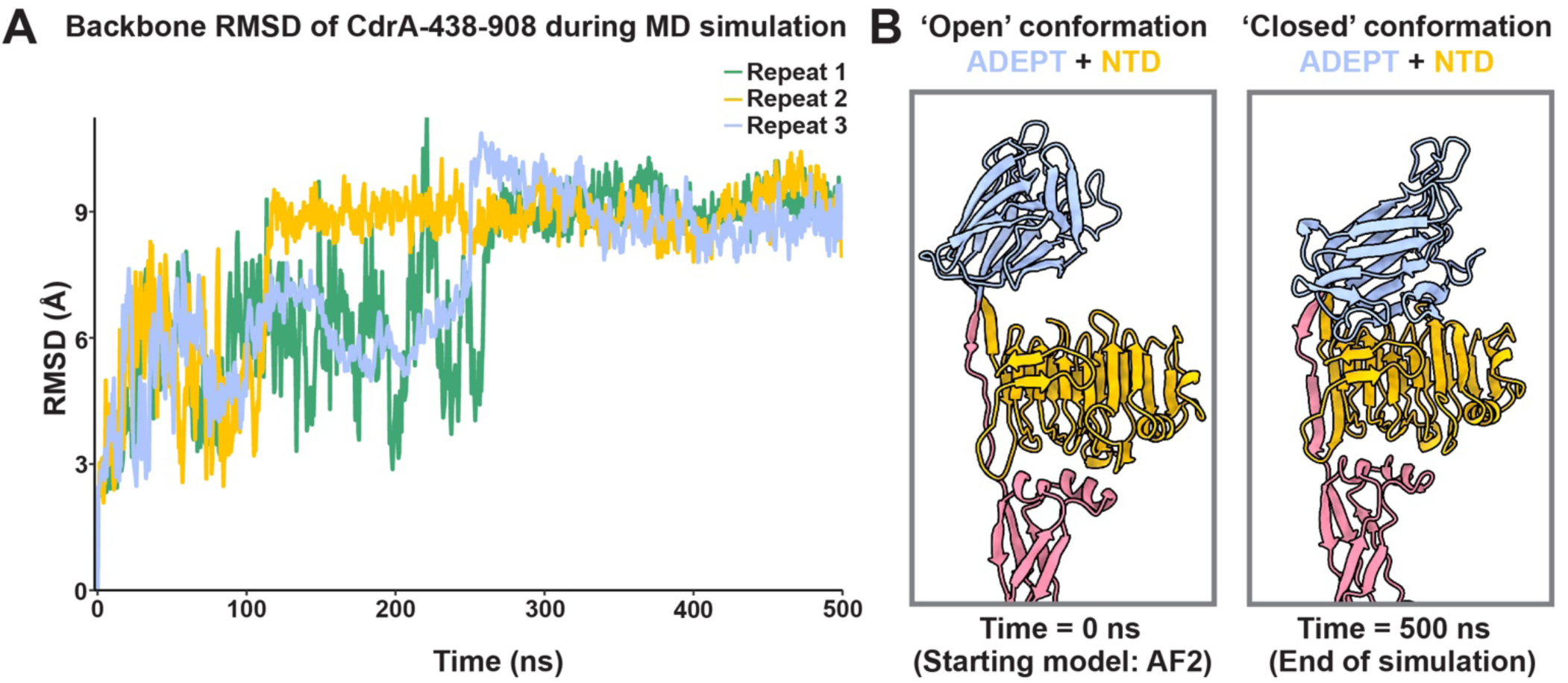
Molecular dynamics simulations of the CdrA N-terminus (residues 438-908). **A)** Plot displaying CdrA backbone root mean square deviation (RMSD) changes as molecular dynamics (MD) simulation proceeds. The AlphaFold2 model of the CdrA N-terminus (from residues 438-908 – including NTD, ADEPT and one stalk repeat) was allowed to explore RMSD changes in a solvent box for a total simulation time of 500 ns. The simulation was repeated 3 times, with repeats indicated by differentially coloured lines: repeat 1 = green, repeat 2 = yellow & repeat 3 = blue. **B)** Conformations of CdrA-438-989 at the beginning of the simulation (0 ns) and at the end (500 ns). CdrA-438-989 is coloured by domain, with: NTD = yellow, ADEPT = blue, stalk repeat = pink. At the beginning of the simulation, a much larger distance in the ADEPT:NTD interface is observed with a more ‘open’ conformation. At the end of the simulation, a much closer distance in the ADEPT:NTD interface is observed, with CdrA-438-989 in a ‘closed’ conformation.

**Fig. S5.**
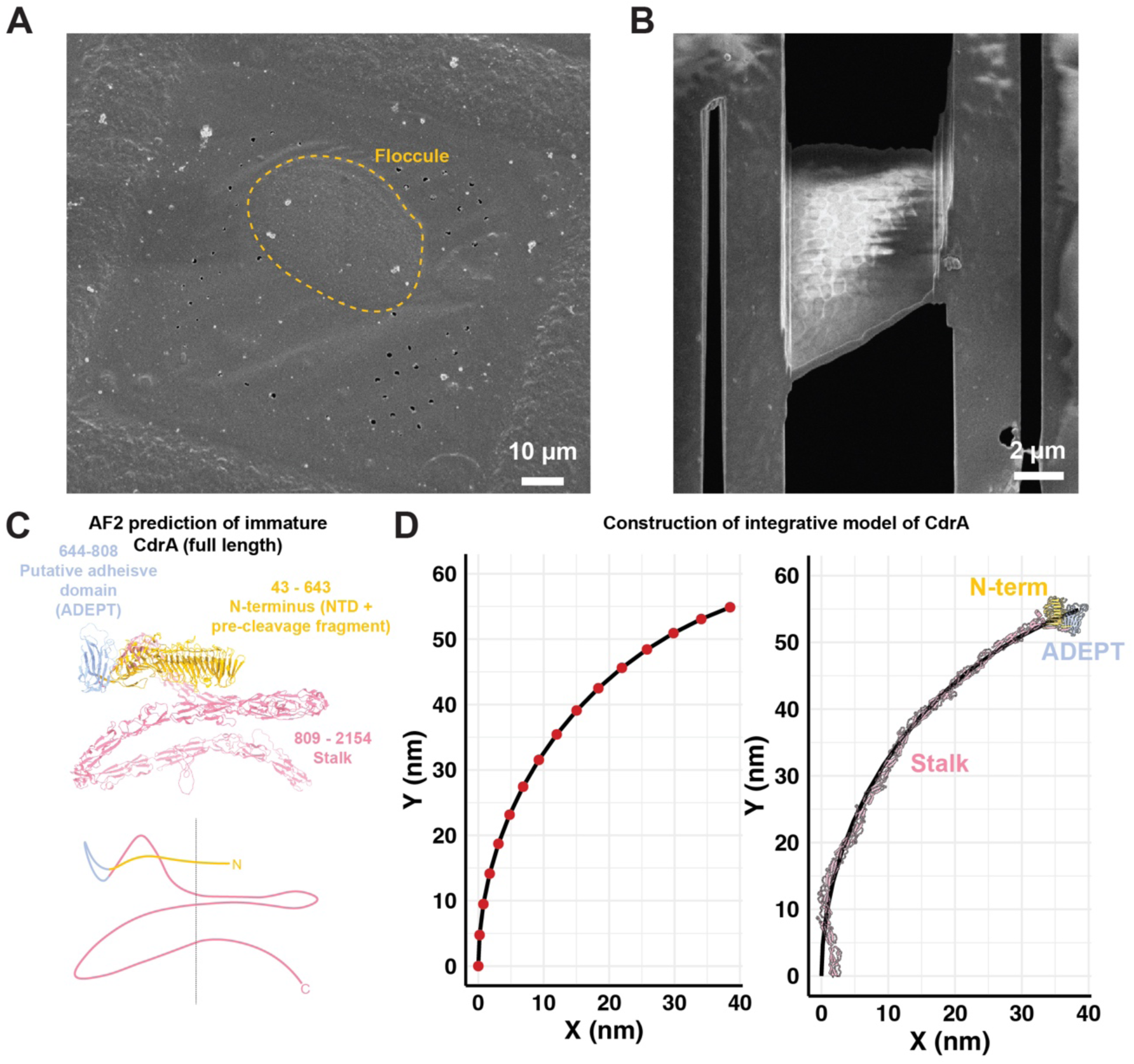
Calculation of the *in situ* model of CdrA. **A)** SEM image of a CdrA-mediated floccule frozen on an electron microscopy grid prior to focused-ion beam (FIB) milling. Region highlighted in yellow corresponds to a CdrA-mediated floccule and was targeted for FIB milling. **B)** SEM image of lamella of floccule produced from FIB milling. **C)** AlphaFold3 prediction of full length CdrA in a condensed, coiled-up architecture. Domain boundaries for full-length CdrA are indicated and the prediction is coloured by domain: N-terminus (NTD + pre-cleavage fragment (residues 43-437)), ADEPT and stalk. **D)** Left hand plot displays an estimation for the curvature of CdrA which was used for production of the *in situ* integrative model of CdrA. The coordinates were used as restraints for molecular dynamics simulation. Coordinates were derived from fitting the average length of CdrA (69nm) with the measured radius of curvature of CdrA (58 nm; with the assumption that curvature is constant throughout the CdrA molecule). On the right, the integrative model of CdrA is overlaid onto these points, displaying the fit of the model to the coordinates after molecular dynamics simulations.

**Fig. S6.**
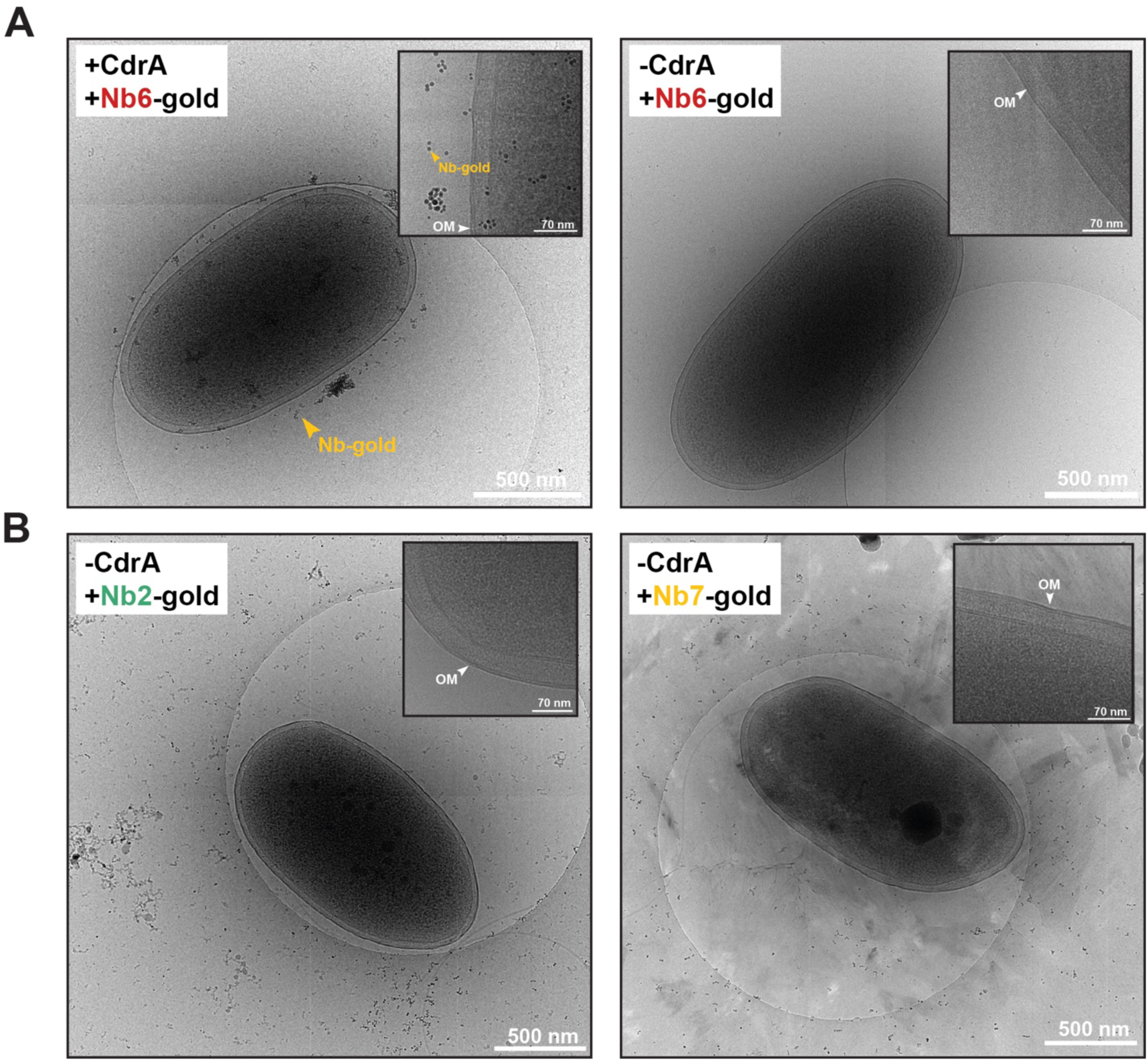
Anti-CdrA nanobodies specifically label CdrA at the cell-surface. **A)** Cryo-EM images of cells labelled with Nb6-gold. Left panel shows a Nb6-gold labelled cell where CdrA is expressed (hence high levels of surface Nb6-gold decoration). The zoomed inset (left) shows that the cell-surface is highly decorated with Nb6-gold. The right image shows an unexpressed CdrA control where no CdrA is expressed and as a result, there is no Nb6-gold labelling at the cell-surface. The zoomed inset (right) also displays an absence of Nb6-gold labelling at the cell-surface. OM = outer membrane. For all images, the scale bar in the main panel represents 500 nm and scale bar in the inset represents 70 nm. **B)** Cryo-EM images of CdrA controls for Nb2-gold (left hand side) and Nb7-gold (right hand side) on cells with no CdrA expression. Scale bars = 500 nm. For both, there is also no cell-surface decoration of Nb-gold (see insets). In the right-hand main image for Nb7-gold, the features around the cell are freezing artefacts due to crystalline ice. For all images, the scale bar in main panel represents 500 nm and the scale bar in inset represents 70 nm.

**Fig. S7.**
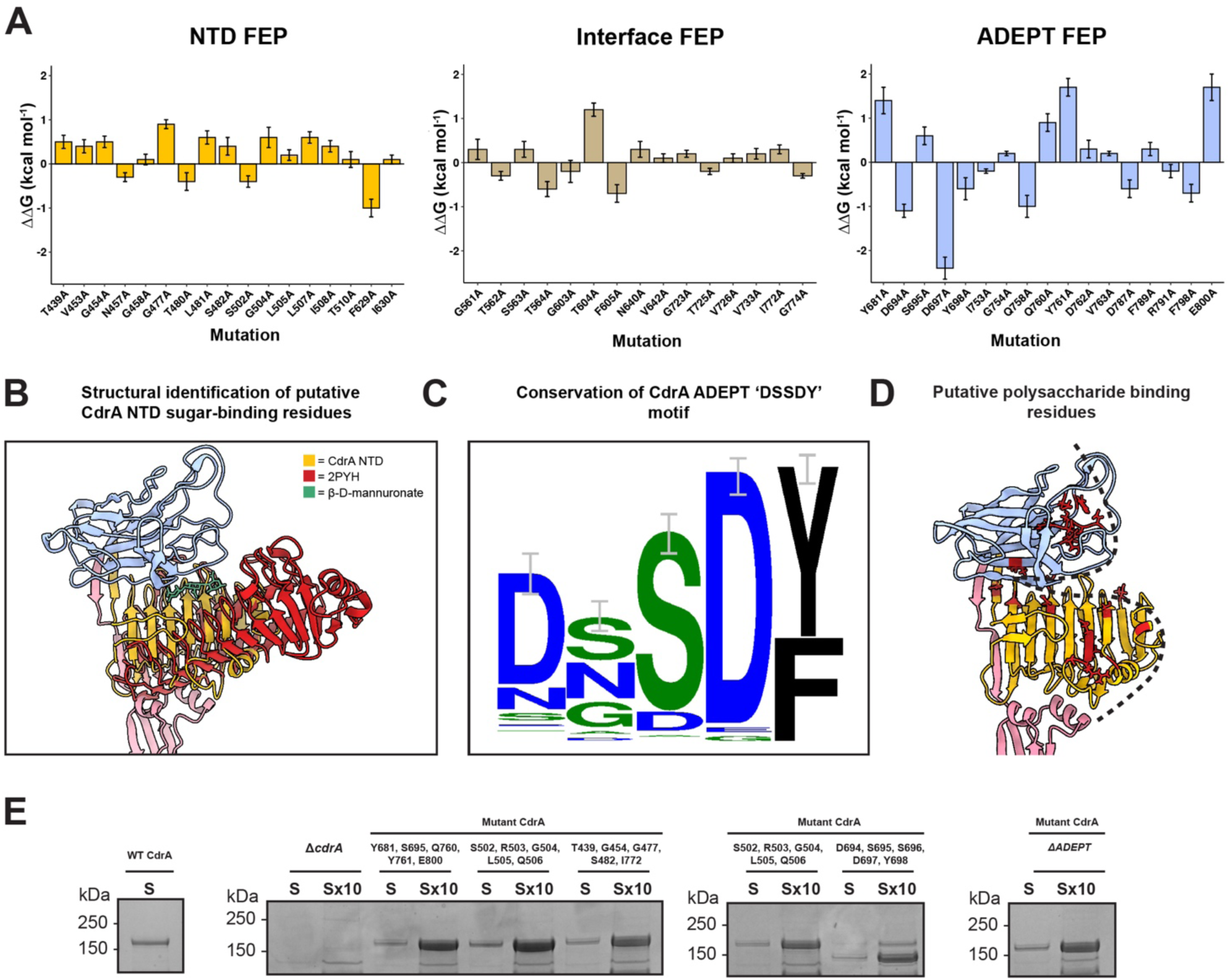
Mutations to CdrA to probe sugar binding. **A)** Free energy perturbation (FEP) plots displaying the change in free energy (ΔΔG) when the specified residues were mutated to alanine. A positive ΔΔG value suggests that mutation to alanine decreases the affinity to Psl compared to the WT. Plots shown are for residues in the NTD (yellow), the ADEPT:NTD interface (beige) and the ADEPT (blue). **B)** Superposition of the cryoEM structure (from this study) of the CdrA NTD (yellow) with a crystal structure of the right-handed β-helix domain of mannuronan C-5 epimerase AlgE4 – PDB code: 2PYH (red). The position of the co-crystallised sugar β-D-mannuronate (green) indicated putative sugar binding residues in the NTD that were mutated. **C)** Sequence alignment of the ‘DSSDY’ motif in the loop region of the CdrA ADEPT shows a high level of sequence conservation. Error bars represent the approximate Bayesian 95% confidence interval. The ‘DSSDY’ motif logo shown here was produced using Weblogo (Crooks *et al*., 2004). **D)** Mapping of all the experimentally mutated residues to the CdrA adhesive N-terminus reveals a putative polysaccharide binding pocket (indicated by dashed black line). **E)** SDS-PAGE analysis of sheared protein from the surface of cells overexpressing mutant CdrA constructs. Molecular weight standards are indicated on the left. CdrA expected molecular weight is ∼176 kDa. CdrA Δ*ADEPT* expected molecular weight is 156 kDa. S = total sheared protein; Sx10 = total sheared protein 10x concentrated.

## Tables

**Table S1.**
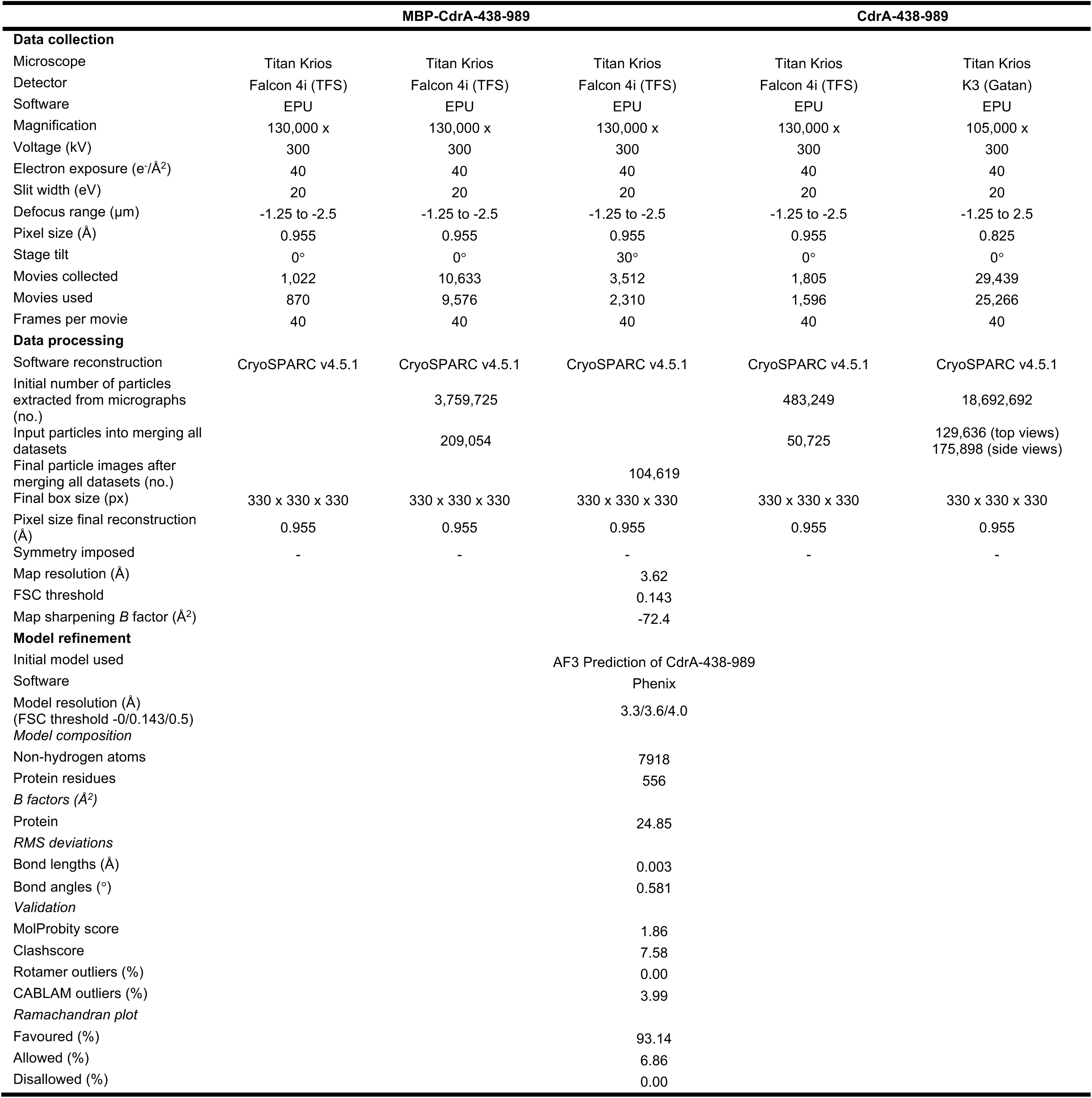
– Cryo-EM data collection, refinement, and validation statistics.

|  | MBP-CdrA-438-989 |  |  | CdrA-438-989 |  |
| --- | --- | --- | --- | --- | --- |
| Data collection |  |  |  |  |  |
| Microscope | Titan Krios | Titan Krios | Titan Krios | Titan Krios | Titan Krios |
| Detector | Falcon 4i (TFS) | Falcon 4i (TFS) | Falcon 4i (TFS) | Falcon 4i (TFS) | K3 (Gatan) |
| Software | EPU | EPU | EPU | EPU | EPU |
| Magnification | 130,000 x | 130,000 x | 130,000 x | 130,000 x | 105,000 x |
| Voltage (kV) | 300 | 300 | 300 | 300 | 300 |
| Electron exposure (e<sup>-</sup>/Å<sup>2</sup>) | 40 | 40 | 40 | 40 | 40 |
| Slit width (eV) | 20 | 20 | 20 | 20 | 20 |
| Defocus range (µm) | -1.25 to -2.5 | -1.25 to -2.5 | -1.25 to -2.5 | -1.25 to -2.5 | -1.25 to 2.5 |
| Pixel size (Å) | 0.955 | 0.955 | 0.955 | 0.955 | 0.825 |
| Stage tilt | 0° | 0° | 30° | 0° | 0° |
| Movies collected | 1,022 | 10,633 | 3,512 | 1,805 | 29,439 |
| Movies used | 870 | 9,576 | 2,310 | 1,596 | 25,266 |
| Frames per movie | 40 | 40 | 40 | 40 | 40 |
| Data processing |  |  |  |  |  |
| Software reconstruction | CryoSPARC v4.5.1 | CryoSPARC v4.5.1 | CryoSPARC v4.5.1 | CryoSPARC v4.5.1 | CryoSPARC v4.5.1 |
| Initial number of particles extracted from micrographs (no.) |  | 3,759,725 |  | 483,249 | 18,692,692 |
| Input particles into merging all datasets |  | 209,054 |  | 50,725 | 129,636 (top views)<br>175,898 (side views) |
| Final particle images after merging all datasets (no.) |  |  | 104,619 |  |  |
| Final box size (px) | 330 x 330 x 330 | 330 x 330 x 330 | 330 x 330 x 330 | 330 x 330 x 330 | 330 x 330 x 330 |
| Pixel size final reconstruction (Å) | 0.955 | 0.955 | 0.955 | 0.955 | 0.955 |
| Symmetry imposed | - | - | - | - | - |
| Map resolution (Å) |  |  | 3.62 |  |  |
| FSC threshold |  |  | 0.143 |  |  |
| Map sharpening *B* factor (Å<sup>2</sup>) |  |  | -72.4 |  |  |
| Model refinement |  |  |  |  |  |
| Initial model used |  |  | AF3 Prediction of CdrA-438-989 |  |  |
| Software |  |  | Phenix |  |  |
| Model resolution (Å)<br>(FSC threshold -0/0.143/0.5) |  |  | 3.3/3.6/4.0 |  |  |
| Model composition |  |  |  |  |  |
| Non-hydrogen atoms |  |  | 7918 |  |  |
| Protein residues |  |  | 556 |  |  |
| *B* factors (Å<sup>2</sup>) |  |  |  |  |  |
| Protein |  |  | 24.85 |  |  |
| RMS deviations |  |  |  |  |  |
| Bond lengths (Å) |  |  | 0.003 |  |  |
| Bond angles (°) |  |  | 0.581 |  |  |
| Validation |  |  |  |  |  |
| MolProbity score |  |  | 1.86 |  |  |
| Clashscore |  |  | 7.58 |  |  |
| Rotamer outliers (%) |  |  | 0.00 |  |  |
| CABLAM outliers (%) |  |  | 3.99 |  |  |
| Ramachandran plot |  |  |  |  |  |
| Favoured (%) |  |  | 93.14 |  |  |
| Allowed (%) |  |  | 6.86 |  |  |
| Disallowed (%) |  |  | 0.00 |  |  |

**Table S2.** Mutations to the ADEPT, ADEPT:NTD interface and NTD. For each penta-alanine mutation set, the domain or interface of CdrA that the mutation set targets is shown as well as the source for each mutation.

| Mutation set | Targeting | Source of mutations |
| --- | --- | --- |
| T439A, G454A, G477A, S482A, I772A | NTD | FEP simulations |
| S502A, R503A, G504A, L505A, E506A | NTD | Structural alignment with PDB:2PYH<br>(same superfamily as CdrA NTD) and<br>subsequent docking with Psl<br>exopolysaccharide |
| Q561A, S563A, T604A, N640A, I772A | ADEPT:NTD | FEP simulations |
| Y681A, S695A, Q760A, Y761A, E800A | ADEPT | FEP simulations |
| D694A, S695A, S696A, D697A, Y698A | ADEPT | Comparison with other ADEPT family<br>members – ‘DSSDY’ motif |

## Movie Legends

**Movie S1. Cryo-EM structure of the CdrA adhesive N-terminus.**

A 3.6 Å resolution cryo-EM density map of the adhesive N-terminus is displayed. This density map was used to build an atomic model of the adhesive N-terminus (including the ADEPT, NTD and two stalk repeats). The map and model are coloured by domain (ADEPT – blue, NTD – yellow, stalk – pink) and both surface depiction and ribbon diagrams are shown.

**Movie S2. *In situ* arrangement of CdrA.**

A tomogram of a FIB-milled CdrA-dependent floccule is displayed which is segmented to display cell membranes (green) and the fibrillar CdrA molecules (pink).

